# Hidden-driver inference reveals synergistic brain-penetrant therapies for medulloblastoma

**DOI:** 10.1101/2025.11.20.689490

**Authors:** Jingjing Liu, Xu Yang, Mingrui Zhu, Xinran Dong, Honglei Zhou, Brandon Bianski, Barbara Jonchere, Wenwei Lin, Xiang Fu, Sheetal Bhatara, Jiyuan Yang, Song-Eun Lim, Lei Yang, Burgess B. Freeman, Abigail S. Wang, Ruilin Jiang, Taosheng Chen, Giles W. Robinson, Martine F. Roussel, Thomas E. Merchant, Amar Gajjar, Jiyang Yu

**Author notes:** These authors contributed equally (Jingjing Liu, Xu Yang, Mingrui Zhu).

## Abstract

Effective therapies for high-risk medulloblastoma (MB), particularly MYC-driven Group 3 (G3) MB, remain elusive due to limited druggable mutations, poor blood–brain barrier (BBB) penetration, and rapid resistance. We developed **SINBA** (**Synergy Inference by Data-driven Network-Based Bayesian Analysis**), a systems biology framework that computationally prioritizes synergistic, BBB-permeable drug combinations by identifying hidden drivers sustaining oncogenic programs. Integrating MB-specific networks, transcriptomic data, and drug–gene interactions, SINBA nominated 32 candidates, of which 19 were experimentally validated as synergistic. Through iterative prioritization and experimental refinement, the **MEK inhibitor mirdametinib** and **p38 inhibitor regorafenib** emerged as the top brain-penetrant pair, suppressing G3 MB progression and extending survival in xenograft and immunocompetent models, with efficacy enhanced by low-dose radiation. Single-cell analysis revealed selective targeting of **developmental origins** and immune reprogramming. These findings establish SINBA as a computationally assisted discovery framework for clinically actionable combinations in high-risk MB.

## INTRODUCTION

Brain tumors are the most common pediatric solid tumor, accounting for more deaths in children under the age of 20 than any other pediatric cancer^1^. Among them, the most prevalent is medulloblastoma (MB), an embryonal tumor of the cerebellum that consists of four distinct molecular subtypes: WNT, SHH, Group 3 (G3), and Group 4 (G4)^2, 3^. Despite the unique molecular and clinical characteristics of these subtypes, standard treatment for medulloblastoma follows a single protocol: maximal surgical resection followed by craniospinal irradiation and adjuvant chemotherapy. While these interventions have improved the 5-year survival rate to over 80% for average-risk patients^4, 5^, one in three high-risk MB patients will relapse within five years, and the survival rate for relapsed patients remains dismal^6, 7^.

Common approaches to therapeutic discovery have yielded few clinically effective treatments for high-risk MB, due to both the nature of the disease and the limitations of current methods^8^. Most notably, few high-risk MB cases possess well-characterized druggable alterations—in fact, approximately 20% lack any identifiable genetic mutations—making the development of genomics-driven targeted therapies difficult^9^. High-throughput screening (HTS), though useful in research contexts, produces single agent therapies that may have poor blood-brain barrier (BBB) permeability, that are susceptible to rapid tumor resistance, and whose efficacy often depends on the relevance of the disease model used and the extent to which it recapitulates patient tumor biology.

The recent therapeutic success of combination therapies in certain cancers has generated widespread interest in drug combination screening^10, 11, 12, 13, 14, 15, 16, 17, 18, 19, 20, 21^; however, standard experimental methods for combination screening are costly and limited in scale. Computational approaches^22, 23, 24, 25, 26, 27^ (such as RECODR^25^ or a graph-embedding machine learning approach of single-cell transcriptomics data, DrugCell^27^ or the community-developed approaches generated by the DREAM Consortium) have been developed to circumvent these limitations, but their utility in G3 MB is limited by, for example, the nature of their training datasets or their pan-cancer design.

Although few clinically actionable genes have been identified for MB^28^, previous studies have identified cancer driver genes that, rather than possess genetic mutations, are functionally dysregulated through epigenetic modifications or post-translational alterations^29, 30, 31, 32, 33^. Such “hidden” drivers could be effective therapeutic targets; however, their lack of genetic mutation makes them more challenging to identify. We therefore developed SINBA, an end-to-end workflow that uses network-based Bayesian analysis to identify putative bottleneck drivers and then, based on those drivers, prioritizes candidate therapeutic drug combinations for experimental refinement. This approach addresses two of the major challenges of therapeutic discovery in MB: identifying druggable targets and reducing the candidate drug pool for combination screening.

In this study, we applied SINBA to G3 medulloblastoma, leveraging transcriptomic data of three medulloblastoma patient cohorts. Through iterative computational prioritization, experimental screening, and pharmacologic optimization, we identified regorafenib and the MEK inhibitor mirdametinib as a promising brain-penetrant combination. We validated the synergistic effects of this drug pair through in vitro models and tested its efficacy in preclinical studies using both human xenografts and immunocompetent syngeneic murine G3 MB models. Combining regorafenib-MEK inhibitor treatment with low-dose irradiation improved cancer cell killing, supporting the potential benefit of incorporating these agents into standard MB therapy. Single-cell RNA sequencing (scRNA-seq) analysis revealed elevated driver activity and enhanced drug sensitivity scores in tumor cells that transcriptomically resemble early Unipolar Brush Cell (UBC) progenitors (henceforth “early UBC progenitor-like cells”), suggesting susceptibility to this combination therapy. Critically, this combination profoundly remodeled the tumor microenvironment (TME) in an immunocompetent syngeneic mouse model, demonstrating both direct and immunomodulatory tumor suppression effects.

## RESULTS

### Nominating synergistic brain-penetrant drug combinations for G3 MB using SINBA, a hidden-driver-based computational framework

SINBA is a network-based algorithm that prioritizes candidate synergistic drug combinations based on subtype-specific drivers. Broadly, the SINBA workflow proceeds as follows: users first leverage a disease-specific gene-gene interaction network to identify drivers upregulated in a selected cancer subtype (candidate drivers are inferred from gene activity and do not imply causality); these drivers are then filtered for druggability and provided as input to the SINBA software. SINBA then uses the interaction network and a drug-target database to design HTS experiments, the results of which are fed back into the software as additional input. Using these data, SINBA matches the input drivers with other hub genes to nominate gene pairs with potential synergistic activity; these driver pairs are translated into drug pairs for subsequent combination screening. Finally, the results of combination screening are analyzed for synergy using a third-party algorithm, SynergyFinder^34, 35^; the drug pairs are then ranked according to their synergy. It should be noted that driver selection and driver-drug pairing are interdependent, concurrent processes. For example, the input drivers are used to select drugs for HTS, but then the results of HTS are used to further refine the list of drivers.

Using the G3 subtype of MB as a proof-of-concept, we describe the SINBA workflow in greater detail below (**Fig. 1a**):

**Figure 1.**
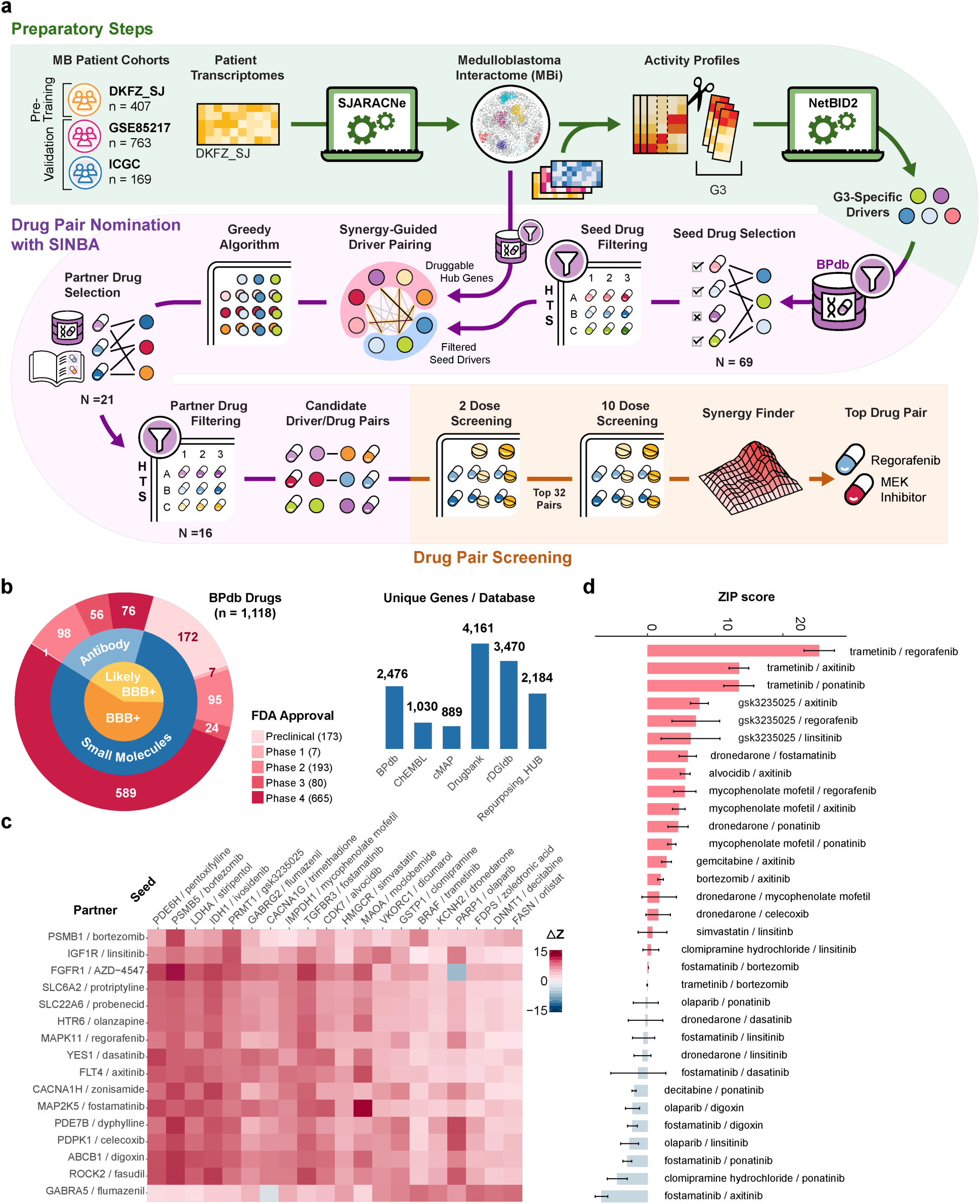
SINBA leverages gene activity data and knowledge-based drug databases to predict and guide screening of synergistic drug pairs for high-risk medulloblastoma. **a** Overview of the SINBA workflow for identifying synergistic drug combinations in medulloblastoma. Preparatory steps include using SJARACNe to generate a disease-specific gene interaction network and NetBID2 to identify subtype-specific driver genes. SINBA then selects drugs for single-agent screening before predicting synergistic drug pairs for combination testing. Finally, combination screening and synergy analysis identify promising drug pairs for validation. **b** Breakdown of the Brain Penetrant Database (BPdb), a manually curated repository of potential and clinically validated blood-brain barrier penetrating (BBB+) drugs. SINBA integrates the BPdb with five additional drug-gene interaction databases for comprehensive target annotation. **c** An optimized driver pair matrix based on SINBA’s predictions of synergy in silico. Seed drivers (columns) and partner drivers (rows) are listed with their respective drugs. Delta Z-scores indicate the predicted synergy of each driver pair. **d** Synergy analysis of the results of 10-dose combination screening. Synergy is expressed as a Zero Interaction Potency (ZIP) score. Data are represented as the mean ± SEM.

#### Preparatory Steps

To equip SINBA to select synergistic drug pairs with good blood-brain barrier permeability, we generated a Brain Penetrant drug-target database (BPdb), comprising 1,118 established or potentially brain-penetrant agents targeting 2,476 genes (**Methods; Fig. 1b**). The BPdb expands substantially on the B3DB^36^, a database of rigorously validated brain-penetrating compounds (**Supplementary table 4**), by including additional, potentially-penetrating agents identified in the literature and clinical trials. Agents were considered as potential penetrants if they had been evaluated in neurotherapeutic trials or represented antibody-based modalities. 41% of the drugs in the BPdb were selected in this manner; overall, 59% of the drugs are FDA-approved. Five drug-gene interaction databases were used to annotate the targets in the BPdb: DrugBank^37^, ChEMBL^38^, cMAP^39^, Repurposing Hub^40^, and DGIdb^41, 42, 43^ (**Fig. 1c; Supplementary Table 1 and 4**). Along with the BPdb, these drug-target databases were integrated into SINBA, supplying it with a comprehensive set of drugs and pharmacologically targetable genes to maximize generalizability outside of central nervous system tumors.

After compiling the BPdb, we pre-trained a disease-specific, gene-gene interaction network for medulloblastoma by applying SJARACNe^44^, a mutual-information-based algorithm, to a dataset of 407 human MB patient transcriptome profiles (DKFZ_SJ)^28, 45, 46^. The resulting medulloblastoma interactome (MBi) captures the transcript-level interactions between transcription factor (TF) and signaling (SIG) genes^47^—the network hubs—and their predicted regulons. Altogether, the interactome comprised 1,567,381 interactions between 38,621 transcripts (corresponding to 15,317 genes); 12,495 of these transcripts belonged to TF and SIG hub genes (7,155 total).

Finally, we used the pretrained MBi and our NetBID2^48^ algorithm to identify G3 MB-specific drivers, henceforth referred to as “seed” drivers (see **Methods** for details). To do so, we superimposed the MBi onto the transcriptomic profiles of patients in the pretraining cohort plus two additional validation cohorts, GSE85217^49^ and ICGC^9^ (**Supplementary Table 2**), generating activity profiles for candidate TF and SIG drivers from the gene expression data. The activity inferred for each candidate driver consisted of the expression of downstream targets weighted by the interaction strength. We then used NetBID2 to analyze the activity of the pretraining cohort, revealing 75 candidate drivers that exhibited significantly higher activity in G3 MB compared to the other MB subgroups and normal cerebellum. Drivers for which there were no targeting drugs in the BPdb were discarded, leaving 45 druggable “seed drivers.”

Comparison of the activity and expression profiles of these seed drivers in the two validation cohorts supports SINBA’s activity-based approach to driver nomination. Seed drivers were found to have consistent differential activity but not expression patterns across cohorts (**Supplementary Fig. 1a**), suggesting that activity data may be a more reliable measure of gene function. Data from the GSE85217 cohort, for which clinical outcomes were available, suggested that driver activity may also have greater clinical relevance. Among the 42 drivers (of the 45 druggable “seed drivers”) with matched activity and expression data in this cohort, patient survival was significantly more strongly associated with driver activity than with driver gene expression. Specifically, 67% of the drivers showed a significant association between activity level and patient survival (P < 0.05), whereas only 26% showed a similar association based on gene expression (**Supplementary Fig. 1b**; **Supplementary Table 3**). For example, the activity but not expression of BRAF and CDK7 significantly correlated with overall patient survival (**Supplementary Fig. 1c**). Compared to their RNA expression levels, these drivers’ inferred activities had a cleaner pattern of elevation in G3 MB compared to the other medulloblastoma subtypes. Likewise, in an independent proteomics cohort of MB patient samples^50, 51^, there was an elevation in the phosphorylated protein activity of these genes despite their total protein levels remaining unchanged (**Supplementary Fig. 1d and 1e; Supplementary Table 3**).

Similarly, expression of MYC—whose activation is the hallmark of G3 MB—was insufficient to stratify risk groups in the GSE85217 cohort; on the other hand, there was a clear delineation in MYC *activity* between high- and low-risk MB (**Supplementary Fig. 1f**). Consistent with the findings for BRAF and CDK7, MYC activity—but not total protein expression—was positively correlated with phospho-MYC levels in G3 MB tumors^50^ (**Supplementary Fig. 1g**). These results demonstrate how activity-based approaches can be used to identify clinically relevant drivers that may evade discovery by conventional RNA and total protein expression analyses. To explore the drug databases, gene expression and patient survival data, and driver activities imputed in this study, we developed an interactive web application at https://yulab-stjude.shinyapps.io/SINBA_MB.

#### Nominating Synergistic Drug Pairs with SINBA Algorithm

Using the 45 seed drivers identified for G3 MB as input, SINBA nominated candidate drug pairs for G3 MB through the following iterative steps:

First, the software selected 69 drugs from the BPdb to undergo HTS, ensuring that each of the 45 seed drivers was a target gene for at least one drug. SINBA then used the results of screening these drugs in a G3 cell line (HDMB03) to select the top 20 most effective drugs with unique seed driver targets.

To identify potential synergistic partner genes for the 20 seed drivers, SINBA generated a list of candidate genes. It did this by filtering the hub genes from the MBi for druggability using the BPdb, yielding approximately 800 candidate “partner drivers”. The candidate set was further refined by eliminating drivers that failed to demonstrate adequate potential synergy with any of the seeds. Candidate drivers were considered for further prioritization if their combined subnetwork with a seed driver yielded higher differential activity than either subnetwork alone. Driver synergy was measured using a custom statistical method that assigns a quantitative “synergy score” to driver pairs (see **Methods**). To select the final partners, SINBA used a genetic algorithm^52^–based optimization approach to identify the 20 × 16 seed-driver/partner-driver matrix that maximized the cumulative synergy score (**Fig. 1c; Methods**).

Next, SINBA selected a different drug for each of the 16 partner drivers, with preference given to drugs from the original library. If no suitable drug could be found in the original library, alternative drugs were selected from the BPdb; these drugs (21 in total) were then screened in the G3 cell line and the results used to select final partners for the remaining drivers (For a summary of the HTS results of our combined libraries (90 drugs total), see **Supplementary Fig. 2a and 2b; Supplementary Table 5**). These 16 final partner drugs were then arranged in a matrix with the 20 seed drugs, producing 320 unique drug pairs for subsequent combination screening (**Fig. 1c**). The corresponding 320 driver pairs exhibited substantially lower activity in normal samples and elevated activity in the G3 subgroup (**Supplementary Fig. 2c**). Through this iterative process of computational prioritization and experimental refinement, SINBA generated a set of candidate drug combinations for further validation.

**Figure 2.**
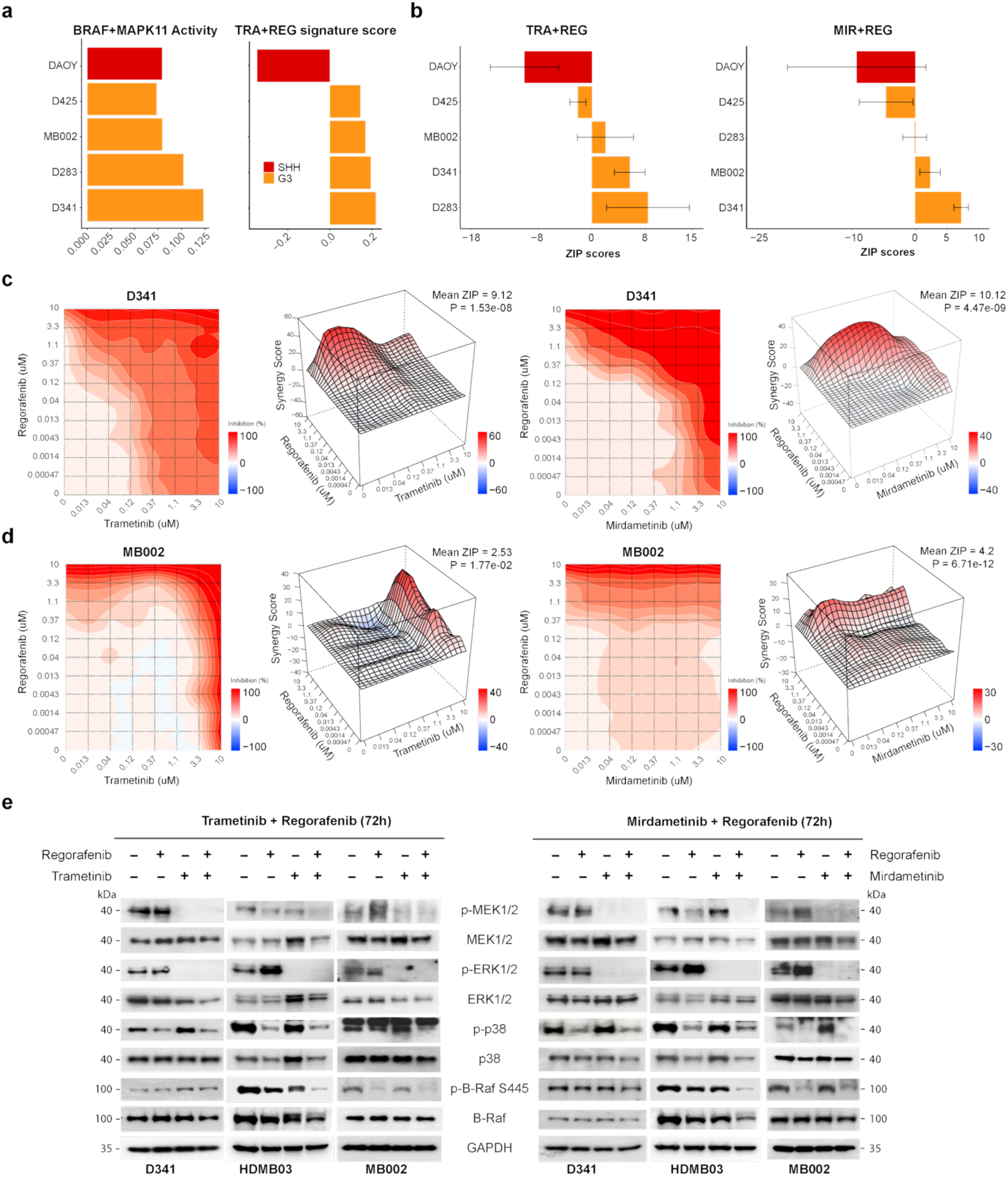
Synergistic and on-target effects of MEK inhibitor and regorafenib combination in human G3 MB models. **a** Left panel shows the combined activities of BRAF and MAPK11 in five human MB cell lines. Right panel shows those cell lines’ drug sensitivity scores for trametinib (TRA) plus regorafenib (REG). **b** Ranked ZIP scores for two regorafenib + MEK inhibitor (trametinib or mirdametinib) combinations in human MB cell lines. Data are represented as mean ± SEM. **c** Cell growth inhibition contour plots and synergy landscapes for drug-dose combinations of regorafenib and trametinib (left) and regorafenib and mirdametinib (right) in D341 cells. **d** Cell growth inhibition contour plots and synergy landscapes for drug-dose combinations of regorafenib and trametinib (left) and regorafenib and mirdametinib (right) in MB002 cells. **e** Western blot analysis comparing total and phosphorylated MEK, ERK, p38, and BRAF following single agent and combination treatment with regorafenib + trametinib (left) or regorafenib + mirdametinib (right) at indicated time points.

#### Drug Combination Screening and Synergy Analysis

Before proceeding to comprehensive drug combination screening, we assessed the potential synergy of our 320 drug pairs through a preliminary two-dose screening. Drugs were tested at EC_20_ (low dose) and EC_50_ (high dose) concentrations; drugs lacking EC_20_ or EC_50_ values were assigned the default values of 1 μM and 6 μM, respectively. This dual-dose strategy produced 1,280 unique drug-dose combinations for testing in the HDMB03 model. The efficacy of each drug-dose combination was quantified using a Δinhibition_mean_ metric: the cell growth inhibition of a two-agent combination minus the mean of the inhibitions produced by using each agent alone at the same dose. Remarkably, 95% of the combinations tested produced efficacies greater than the mean of their parts (**Supplementary Fig. 2d**; **Supplementary Table 6**; see **Methods**).

For each drug-dose combination, we also calculated the Δinhibition_max_, the difference in inhibition between the combination and whichever of its constituents produced greater inhibition alone. We then used this metric to select 32 candidate pairs that met one of the following four conditions: At the low-low dose, (1) combined inhibition >90% and Δinhibition_max_ >10%, (2) combined inhibition >50% and Δinhibition_max_ >15%, or (3) combined inhibition >80% and each single agent’s inhibition >70%; or, at any other dose-combination, (4) combined inhibition >50% and Δinhibition_max_ >20%. These 32 drug pairs were advanced to the comprehensive screening step.

Drug pairs were comprehensively screened in HDMB03 cells using a 10 × 10 dose-response matrix, yielding 100 drug-dose combinations per pair. Screening was performed in triplicate. For each replicate matrix, we evaluated the synergy of each drug-dose combination using the Zero Interaction Potency (ZIP) model, a hybrid of the Loewe additivity and Bliss independence models^34, 35^; these values were then averaged to produce an overall ZIP score for the matrix. Each unique drug pair was assigned a ZIP score equal to the average of its three replicate matrices. Notably, 19 of the 32 drug pairs (59.4%) were found to be synergistic (ZIP score >0, **Fig. 1d**; **Supplementary Table 7**), supporting the utility of SINBA as a prioritization tool for identifying synergistic therapeutic candidates. Notably, synergistic effects were evaluated using a standardized dose-range design. Although this simplified screening strategy balanced the technical limitations of our in-house high-throughput screening platform with the goal of identifying clinically translatable drug combinations, optimization of dose ranges based on the EC_50_ values of individual compounds may further improve the detection of synergistic drug pairs in future studies.

Trametinib, a MEK inhibitor, and regorafenib, a multi-kinase inhibitor, emerged as the most synergistic drug pair among those tested. These drugs demonstrated profound synergy not only overall, but also in the low dose regions of their synergy score landscapes (**Supplementary Fig. 2e**).

#### Fine-tuning the drug pair for optimal BBB-penetrability

To confirm the blood-brain-barrier permeability of these drugs, we conducted preclinical pharmacokinetic studies for regorafenib in female CD-1 nude mice (see **Methods**). Following a single oral dose of regorafenib (10 mg/kg), we observed consistent plasma and brain concentrations, with inter- and intra-mouse coefficients of variation ranging from 7.7% to 28.6% (**Supplementary Fig. 3a**). Brain exposure was assessed by determining the unbound brain-to-plasma partition ratio (*K*_p,uu_), a critical parameter in evaluating CNS drug penetration and distribution. Regorafenib demonstrated reasonable brain distribution (K_p,uu_=0.143), which was consistent with prior murine studies, though slightly lower than previously reported 24-hour estimates^53^. Notably, regorafenib has also been shown to have a robust CNS exposure profile^54,55^. Similar methods^56^ found trametinib to have a less favorable brain distribution (K_p,uu_=0.069, **Supplementary Fig. 3b**). We therefore decided to include mirdametinib, another MEK inhibitor whose BBB-permeability has been established in the literature^57, 58^, in our testing. Consistent with existing studies, we found mirdametinib to have a superior K_p,uu_ of 0.243 (**Supplementary Fig. 3c**). Having confirmed regorafinib’s BBB permeability and selected an additional MEK candidate, we advanced the regorafenib-MEK inhibitor combination to preclinical validation as our lead candidate.

### Regorafenib-MEK Inhibitor synergistic combination selectively kills G3 MB cells *in vitro*

Existing chemoproteomic profiling data has shown overlap between MEK inhibitors’ targets and regorafenib’s targets on the MAPK signaling pathway^59^ (**Supplementary Fig. 3d**). This pathway (KEGG MAPK signaling^60^) is significantly upregulated in G3 tumors compared to normal cerebellum (**Supplementary Fig. 3e**), suggesting selective vulnerability in the G3 subtype. To better understand the mechanism underlying the synergy of MEK inhibitors and regorafenib, we expanded our testing to include an additional four G3 MB cell line models (D341, D425, D283, and MB002) as well as an SHH-MB cell line (DAOY).

First, we mapped the RNA-seq profiles of these cell lines^61^ onto our MBi to generate paired activity profiles for MAPK11 (a subunit of p38 MAP kinase) and BRAF, the target drivers for regorafenib and trametinib/mirdametinib, respectively (**Fig. 2a**). We then calculated a signature for the trametinib/regorafenib combination using iLINCS^62^ data on the drugs’ gene expression signatures in a CNS-disease context. To do so, we used the activity profiles previously generated for each cell line to rank the drivers in each line by activity score. Using these rankings as a reference, we then calculated cell line-specific enrichment scores for the genes in which treatment with regorafenib or trametinib induced up- or down-regulation. These enrichment scores were used to calculate an integrated score for each cell line indicating its predicted relative sensitivity to a trametinib-regorafenib combination (**Fig. 2a and Supplementary Fig. 4a**; see **Methods**).

These sensitivity predictions were largely corroborated in *in vitro* testing, where we used AlamarBlue cell viability assays to evaluate the regorafenib-trametinib and regorafenib-mirdametinib combinations in the MB cell lines. For each cell line, 80 drug-dose combinations were tested per drug pair (eight concentrations of MEK inhibitor and ten concentrations of regorafenib); the results were then used to calculate an overall ZIP score for each drug pair as previously.

Overall, the greatest collective synergy was observed in the D341 cell line, which ranked No. 1 for regorafenib/mirametinib and No. 2 for regorafenib/trametinib (**Fig. 2b and 2c**). Reasonable synergistic effects were also observed for both combinations in MB002 (**Fig. 2b and 2d**). As expected, no synergy was observed for these combinations in the SHH-MB line DAOY (**Supplementary Fig. 4b**). Despite being a G3 MB model, D425 was minimally responsive to both drug pairs, consistent with the low synergy score we had predicted for that cell line (**Supplementary Fig. 4c**). Some difference was observed in the performance of the drug pairs within a single cell line. For instance, the trametinib combination produced the highest synergy in D283, consistent with our predictions, whereas the mirdametinib combination produced no synergy in that line (**Fig. 2b and Supplementary Fig. 4d**).

To confirm the mechanism of action of regorafenib and the MEK inhibitors in the context of combination therapy, we compared the effects of combination therapy vs. monotherapy on signaling in three responsive G3 MB cell lines (D341, HDMB03, and MB002). For Western blot analysis, we selected three well-established targets of MEK inhibition (MEK, ERK, and BRAF), while for regorafenib—a multi-kinase inhibitor—we chose MAPK11/p38, regorafenib’s main target in G3 MB according to our previous analysis with SINBA. Western blot analysis showed that the regorafenib-MEK inhibitor combinations reduced the phosphorylation levels of all four targets (MEK, ERK, BRAF, and p38), while total protein levels remained unchanged. In some cases, combination therapy even produced a greater reduction in a target’s phosphorylation compared to monotherapy. These findings confirmed effective on-target suppression of BRAF-MEK signaling by trametinib/mirdametinib and p38 signaling by regorafenib in G3 MB tumor cells in the contexts of both mono- and combination therapy (**Fig. 2e**).

Together, these *in vitro* studies confirmed the synergistic killing effect of regorafenib-MEK inhibitor combinations on G3 MB cells, but not SHH-MB cells, consistent with the subtype-specific prioritization from SINBA.

### Regorafenib-MEK inhibitor combinations are effective for G3 MB *in vivo*

Having identified D341 as the cell line predicted to be most sensitive to the regorafenib-MEK inhibitor combination (**Fig. 2a**) and having validated this prediction *in vitro* by demonstrating that regorafenib and MEK inhibitors exhibit the greatest synergy in D341 cells (**Fig. 2b**), we moved to investigate whether our findings would also hold *in vivo* (see **Methods**).

Given mirdametinib’s superior brain penetration (compared to trametinib), as well as its greater synergy with regorafenib in the D341 cell line, we chose to advance the regorafenib-mirdametinib combination to *in vivo* testing (**Fig. 2b**). CD-1 nude mice orthotopically implanted with D341 cells were treated with regorafenib (10 mg/kg), mirdametinib (5 mg/kg), or their combination (**Fig. 3a**). Tumor progression was monitored weekly using IVIS imaging beginning one week post-implantation. The combination therapy outperformed both the vehicle control and monotherapy groups in suppressing tumor growth (**Fig. 3b** and **3c**), improving survival (**Fig. 3d**), and increasing the proportion of apoptotic cells (**Fig. 3e**).

**Figure 3.**
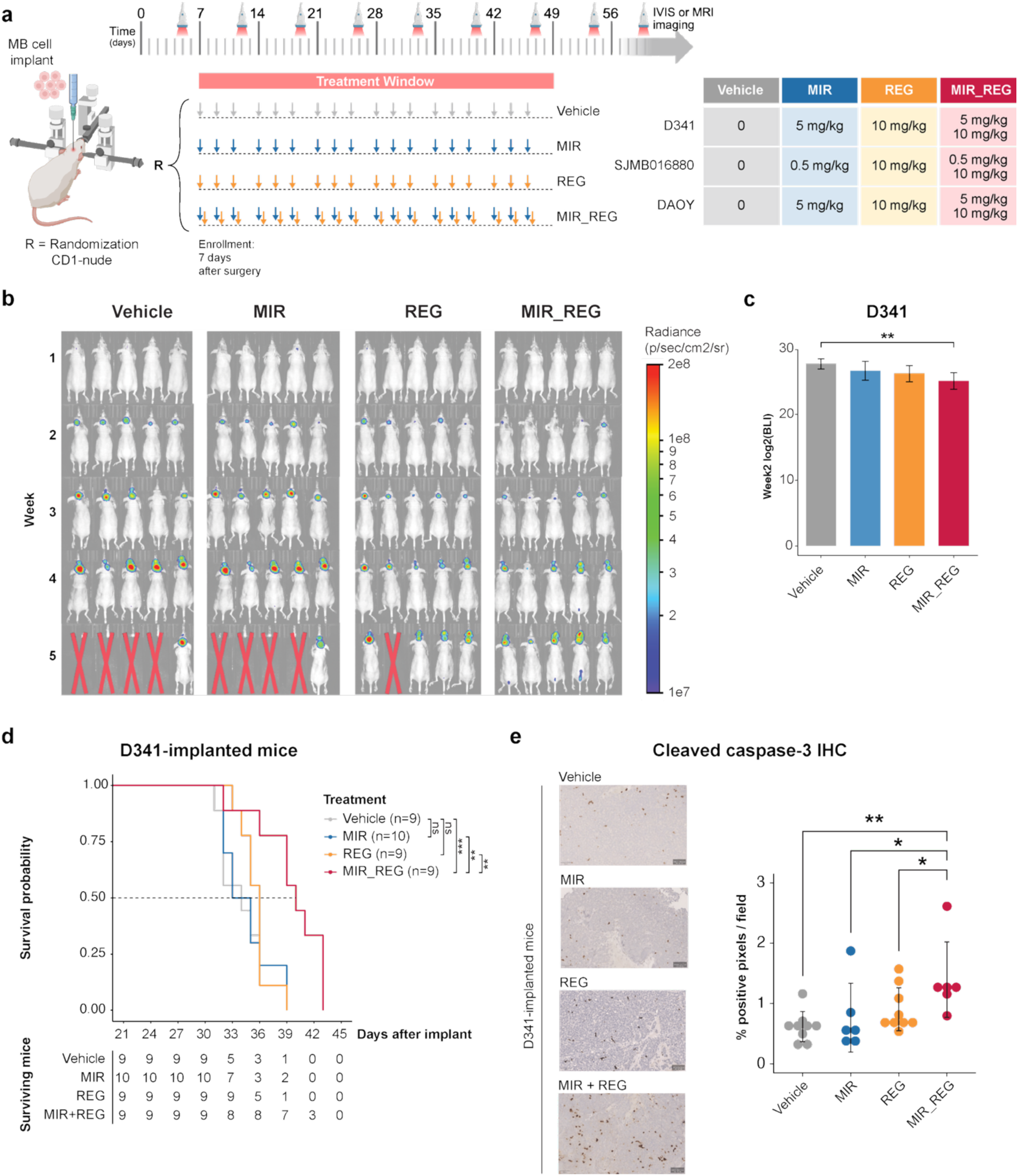
MEK inhibitor and regorafenib combination exhibit synergistic anti-tumor efficacy *in vivo* with G3 cell line model. **a** Schematic of experimental design for drug administration and imaging in CD-1 nude mice implanted intracranially with D341, SJMB016880, or DAOY tumor cells (created with https://BioRender.com). Mice were randomized to receive vehicle (p.o.), mirdametinib (5 mg/kg or 0.5 mg/kg, p.o.), regorafenib (10 mg/kg, p.o.), or the combination, starting one week post-implantation and continuing three times per week for six weeks. IVIS or MRI imaging was performed weekly starting one week after surgery. **b** Representative *in vivo* bioluminescent images of mice bearing orthotopic D341 tumors from weeks 1 to 5 post-implantation. **c** Quantification of bioluminescence signal (BLI) at week 2 post-treatment in the D341 model, showing significant tumor growth reduction in the combination group. Data are presented as mean ± SEM. **d** Kaplan–Meier survival curves for mice implanted with D341 cells and treated as described in **b**. Group sizes (n) are indicated in the risk table. Median survival (days) for the vehicle, mirdametinib, regorafenib, and combination groups was 34, 34, 36, and 40, respectively; log-rank test results: *P < 0.05, **P < 0.01, ***P < 0.001; ns, not significant. **e** Left: Representative IHC staining for cleaved caspase-3 in D341 tumors. Right: Quantification of cleaved caspase-3-positive cells. Data are presented as mean ± SEM. *P < 0.05; **P < 0.01.

To assess the generalizability of this approach, we also tested the *in vivo* efficacy of the mono-and combination therapies in the SHH model DAOY (**Fig. 3a** and **Supplementary Fig. 5a**). Unlike the aggressive D341 model, the DAOY model exhibited a longer median survival of 91.5 days in the control group, compared to 34 days in D341. However, neither the single drug nor combination treatments provided significant survival benefits in the DAOY model (**Supplementary Fig. 5a and 5b**), underscoring the specificity of this therapeutic strategy for G3 MB.

Recognizing the limitations of cell lines in fully recapitulating tumor biology, we extended our investigation to MB patient-derived xenografts (PDXs)^63, 64^. G3 MB PDXs produced significantly higher driver pair activity and drug sensitivity scores than normal cerebellum samples. Among the PDX models, SJMB016880 was predicted to be the most sensitive (**Fig. 4a**). Mice orthotopically implanted with the SJMB016880 model were treated with regorafenib and a reduced dose of mirdametinib (0.5 mg/kg) due to the model’s increased MEK inhibitor sensitivity (**Fig. 4b**). Tumor progression was monitored weekly by MRI. As with the D341 model, combination therapy significantly reduced tumor burden in the SJMB016880 model (**Fig. 4d**) and substantially extended survival (**Fig. 4c**). Mice treated with combination therapy had a median survival of 69 days, compared to 55, 55, and 59 days for mice treated with vehicle, mirdametinib, or regorafenib, respectively. We also observed a greater increase in apoptotic cell percentages following combination therapy than with either monotherapy or vehicle control (**Fig. 4e**).

**Figure 4.**
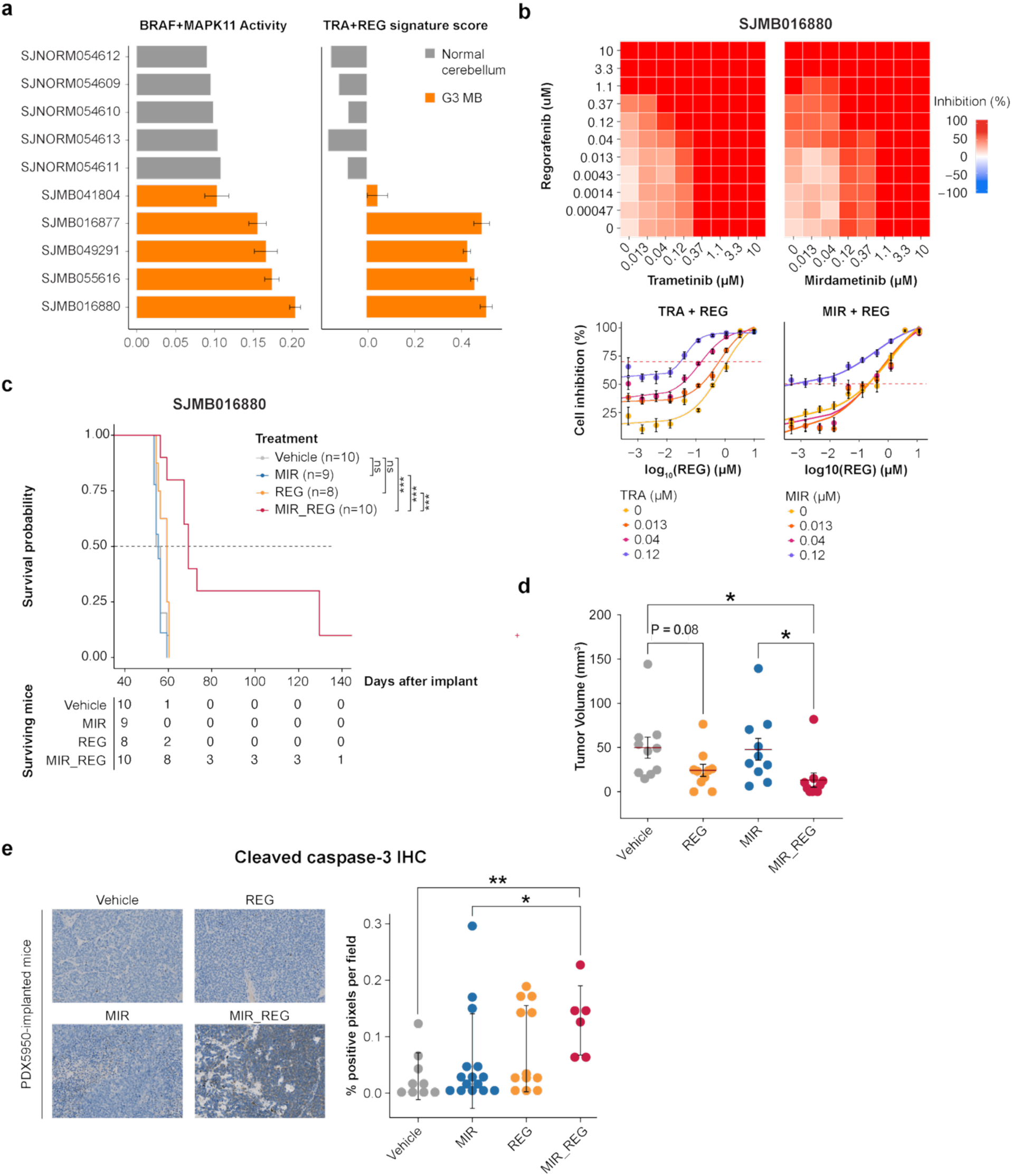
MEK inhibitor and regorafenib combination exhibit synergistic anti-tumor efficacy *in vivo* with patient derived orthotopic xenograft model. **a** Left panel shows the combined activities of BRAF and MAPK11 in normal cerebellum and human G3 MB PDOX models. Right panel shows their drug sensitivity scores for trametinib (TRA) plus regorafenib (REG). **b** Top: Heatmap of SJMB016880 cell viability with regorafenib and MEK inhibitor combinations. Bottom: Dose–response curves for regorafenib in combination with fixed concentrations of trametinib or mirdametinib. **c** Kaplan–Meier survival curves for mice implanted with SJMB016880 and treated as described in Fig. 3a. Group sizes (n) are indicated in the risk table. Median survival (days) for the vehicle, mirdametinib, regorafenib, and combination groups was 55, 55, 59, and 69, respectively; log-rank test results: ***P < 0.001; ns, not significant. **d** MRI-quantified tumor volumes at week 4 post-treatment. *P < 0.05. **e** Left: Representative IHC staining for cleaved caspase-3 in SJMB016880 tumors. Right: Quantification of cleaved caspase-3-positive cells. Data are represented as mean ± SEM. *P < 0.05; **P < 0.01.

**Figure 5.**
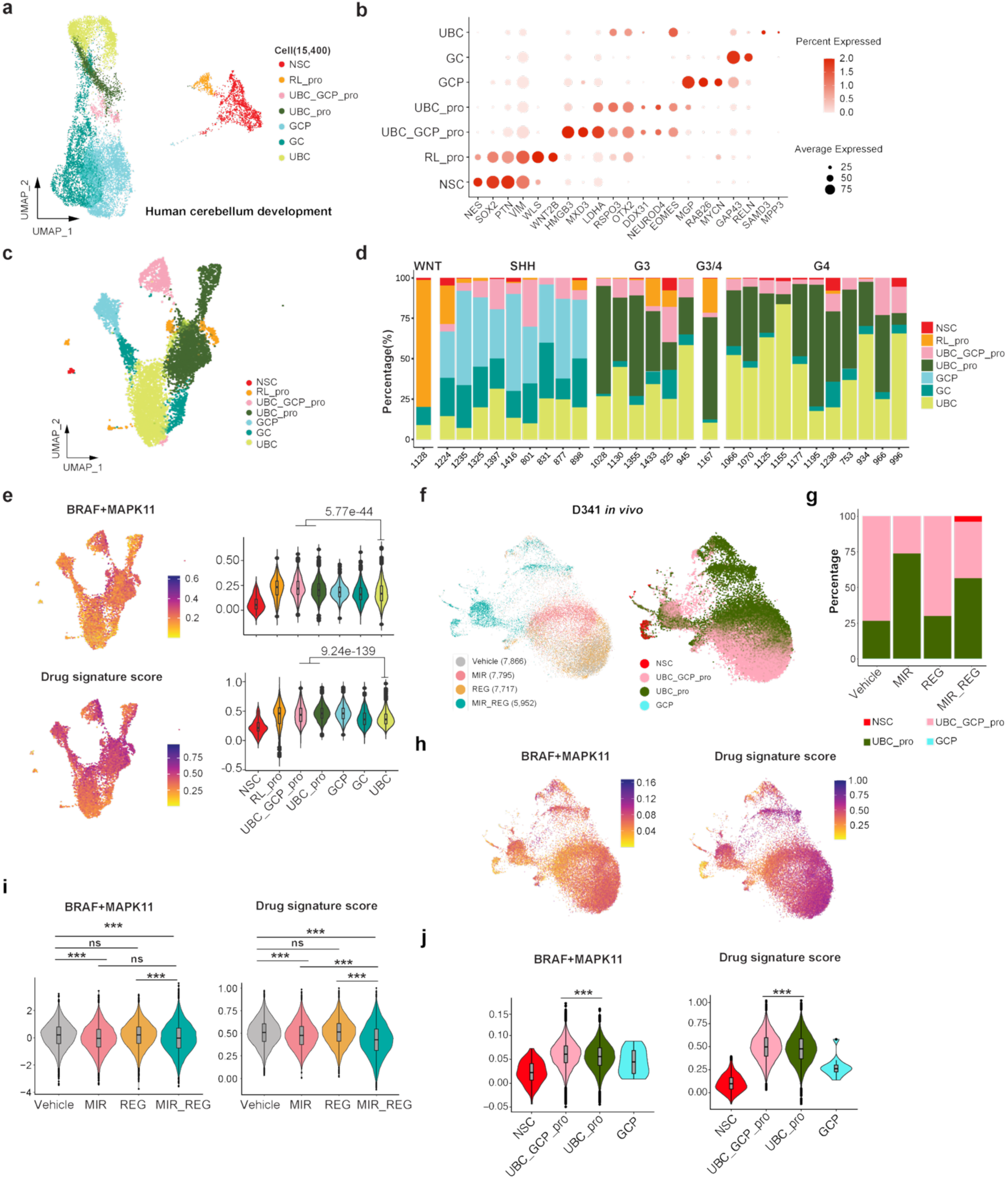
MEK inhibitor and regorafenib combination therapy targets on UBC_GCP common progenitor and UBC progenitor like tumor cells. **a** Re-annotation of the normal glutamatergic neuronal lineage, projected onto a UMAP embedding using scRNA-seq data from Luo et al. **b** Expression patterns of key marker genes defining glutamatergic lineage cell states. **c** Cell state annotation of Medulloblastoma tumor cells (patient scRNA-seq data, GSE156053*; Riemondy et al*.) using the normal developmental reference from A. Visualization restricted to tumor cells. **d** Proportional distribution of tumor cell states across medulloblastoma patient subgroups. **e** Feature and violin plots showing driver combination activities and predicted drug sensitivity scores within tumor populations. **f** UMAP visualization of orthotopically implanted D341 tumor cells (in CD-1 mice) from treated mice, annotated with cell states using the normal developmental reference. **g** Proportional distribution of tumor cell states across treatment groups. **h-j** Single-cell feature plots (**h**) and violin plots (**i**, **j**) of driver activities and drug sensitivity scores across treatment arms (**i**) and predicted cell states (**j**). Two-sided t-tests were used to compare the indicated groups. *P < 0.05, **P < 1.0e-10, ***P < 1.0e-20, ns, not significant.

Body weight loss was noted in the D341-implanted mice three weeks post-implantation regardless of whether they received monotherapy, combination therapy, or vehicle control (**Supplementary Fig. 5c**). Blood analyses showed no significant differences across treatment groups, with all parameters except white blood cell (WBC) count remaining in the normal reference (**Supplementary Fig. 5d**). These findings suggest that weight loss was associated with tumor burden rather than treatment toxicity. In contrast, mice implanted with SJMB016880 experienced slight weight gain during the treatment window. Blood analyses similarly showed no significant differences across treatment groups, though the WBC counts of mice in the single-agent treatment arms fell outside the normal range. Notably, WBC count in the combination group remained within normal limits, suggesting a potential protective effect of the combination therapy (**Supplementary Fig. 5e and 5f**).

### Regorafenib plus MEK inhibitor enhances low-dose radiation therapy *in vivo*

Radiotherapy remains a cornerstone of medulloblastoma treatment^65^, yet patients with G3 MB frequently experience tumor progression and poor survival outcomes^4, 6, 7, 66^. To investigate whether regorafenib and MEK inhibitors could enhance the efficacy of radiotherapy, we tested them in combination with low-dose irradiation using the D341 model. Mice were assigned to four treatment groups: vehicle control, drug combination alone, drug combination administered concurrently with radiation, and drug combination administered following radiation (**Supplementary Fig. 6a**). Radiotherapy was delivered in fractions of 0.5 Gy per day for five consecutive days (see **Methods** for the rationale underlying dose selection). All treatment arms demonstrated improved survival compared with the vehicle control group, with sequential therapy providing the greatest benefit over radiation alone (**Supplementary Fig. 6b and 6c**). Importantly, treatment-related toxicity (measured through body weight and blood analyses) was not observed in any of the experimental groups (**Supplementary Fig. 6d and 6e**).

**Figure 6.**
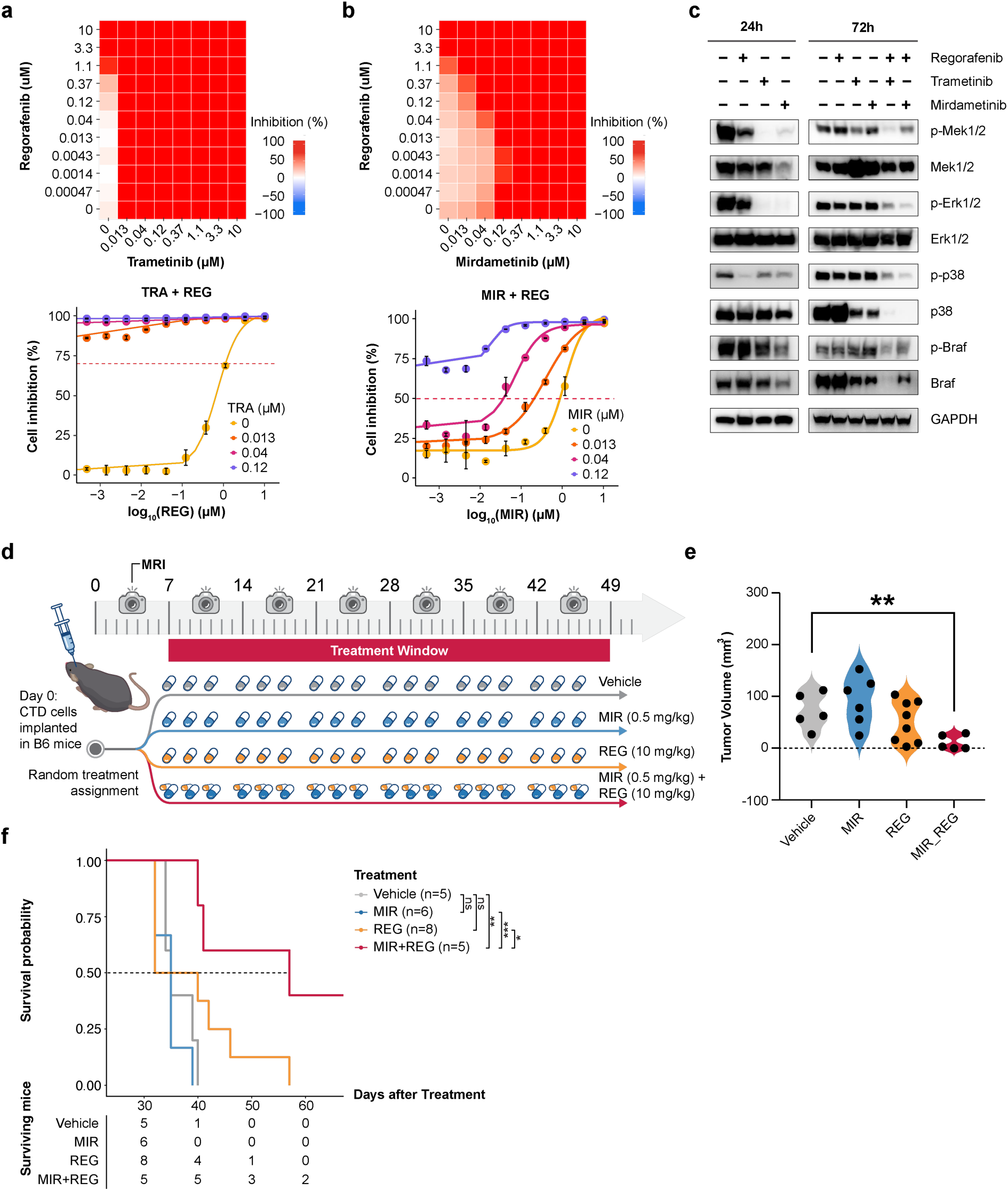
The combination of MEK inhibitor and regorafenib is effective in CTD, an immunocompetent syngeneic mouse model of G3 MB. a-b. Top: Heatmap of the CTD syngeneic mouse tumor cell viability treated with regorafenib and MEK inhibitor combinations. Bottom: Dose-response curves of regorafenib with fixed concentrations of trametinib (left) or mirdametinib (right). **c** Western blot analysis of total and phosphorylated MEK, ERK, p38, and BRAF in CTD cells following treatment at indicated time points. **d** Schematic of experimental design for drug administration and MRI imaging in C57BL/6J mice. Seven days after intracranial implantation of CTD cells, mice were randomized into four treatment groups and treated three times per week for six weeks with vehicle (p.o.), mirdametinib (0.5 mg/kg p.o.), regorafenib (10 mg/kg p.o.), or the combination. **e** MRI-measured tumor volumes across treatment groups. Unpaired two-sided t tests were used; **P < 0.01. **f** Kaplan–Meier survival curves for mice treated as described in (D). Group sizes (n) are indicated in the risk table. Median survival (days) for the vehicle, mirdametinib, regorafenib, and combination groups was 35, 35, 36, and 57, respectively; log-rank test results: *P < 0.05, **P < 0.01, ***P < 0.001, ns, not significant.

Our findings suggest that combining mirdametinib and regorafenib with a markedly reduced radiation regimen (>10-fold lower than conventional clinical doses) improves survival in mice, highlighting the potential of regorafenib–MEK inhibitor combination therapy as a treatment strategy for high-risk G3 medulloblastoma. Further preclinical studies and clinical validation will be necessary to establish optimal radiation dosing and assess the clinical applicability of this approach.

### Single-cell analysis reveals that regorafenib and MEK inhibitors target the developmental origins of G3 MB

Recent single-cell studies of human medulloblastoma have demonstrated significant intra- and inter-tumoral heterogeneity and diverse developmental origins of MB subtypes^67, 68, 69, 70, 71, 72, 73^. To understand how regorafenib-MEK inhibitor combinations affect different cell subpopulations in heterogeneous G3 MB, we analyzed drug target activity using single-cell RNA sequencing (scRNA-seq) data from human developing cerebellum, MB patient tumor samples, the D341 model, and the SJMB016880 xenograft model, both before and after treatment.

To investigate the developmental origins of each MB subtype, we used publicly available scRNA-seq data^71^ to construct a developmental reference for human cerebellum, focusing on the glutamatergic lineage, from which most MB subtypes are thought to arise. Our clustering analysis identified seven distinct developmental states based on previously published markers^74, 75, 76^: neural stem cells (NSC), rhombic lip progenitors (RL_pro), unipolar brush cell and granule cell precursor common progenitors (UBC_GCP_pro), unipolar brush cell progenitors (UBC_pro), granule cells (GC), and unipolar brush cells (UBC) (**Fig. 5a and 5b**; **Supplementary Table 8**). Pseudotime analysis using Slingshot and Monocle 3 reconstructed a putative developmental trajectory in which NSCs first differentiate into RL_pro cells and subsequently into UBC_GCP_pro cells, after which the lineage bifurcates into the UBC and GC branches **(Supplementary Fig. 7a and 7b**; see **Methods**). This single-cell reference for human glutamatergic development is available via an interactive web portal (https://scminer.stjude.org/study/Glutamatergic-lineage).

**Figure 7.**
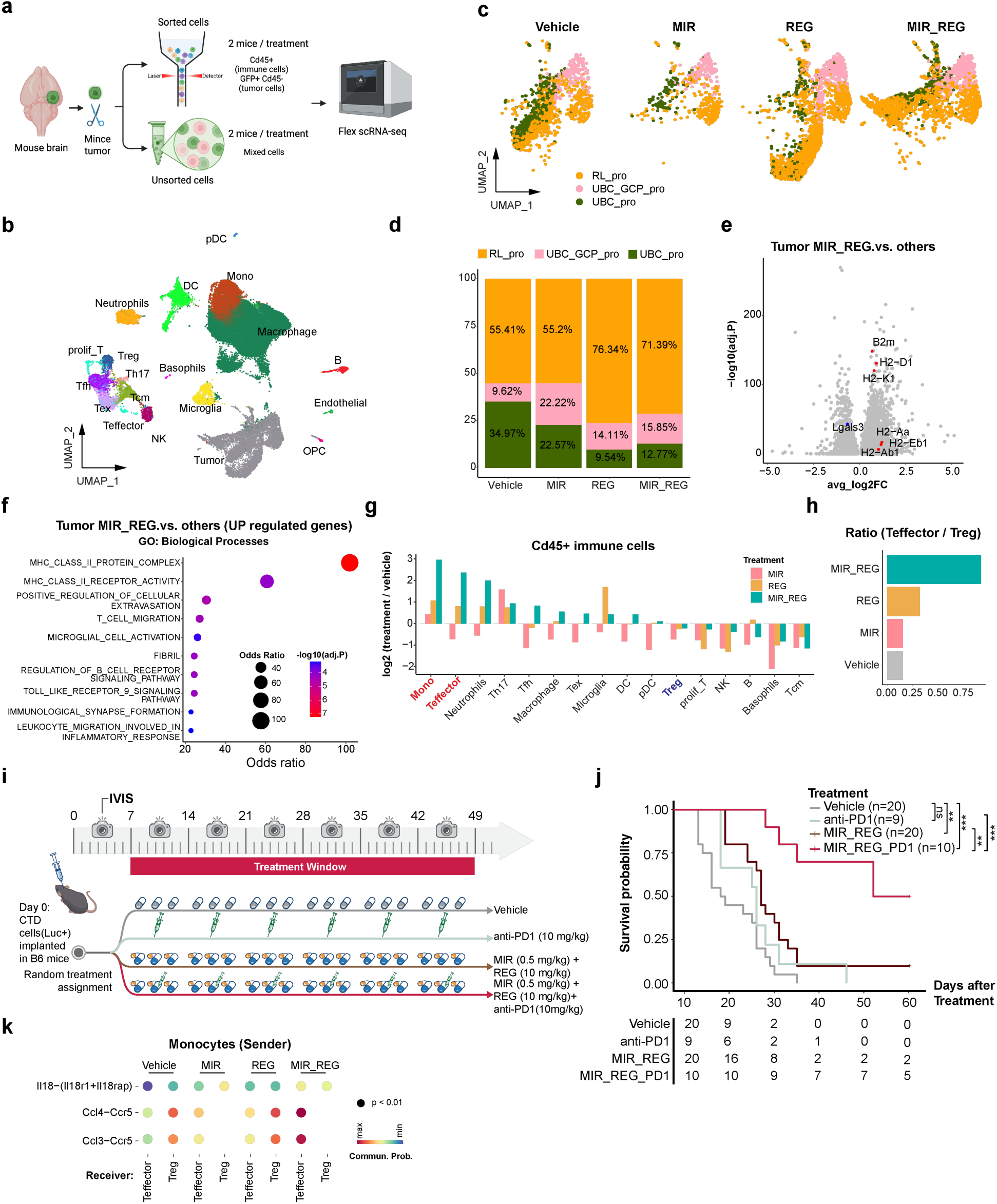
MEK inhibition combined with regorafenib remodels tumor cell states and the tumor microenvironment in the CTD syngeneic mouse model. **a** Schematic of experimental design for generating scRNA-seq data from the CTD syngeneic model preclinical study (created with https://BioRender.com). **b** UMAP visualization of the cells from the CTD model treated with monotherapy or combination therapy *in vivo*, with cells colored by their annotated cell types. **c** UMAP plot of tumor cell states in the four treatment groups. **d** Proportional distribution of tumor cell states across treatment groups. **e** Volcano plot of differentially expressed genes in tumor cells from the MIR_REG combination group versus other treatments. **f** Functional enrichment of top upregulated genes (avg_log2FC > 1, adjusted p < 0.05) in tumor cells from the MIR_REG combination group. **g** Log2 fold-change in abundance of immune cell populations for each treatment group relative to the vehicle control. **h** Ratio of T effector (Teffector) to T regulatory (Treg) cells across treatment groups. **i** Experimental design for evaluation of anti-PD-1 combination therapy. C57BL/6J mice bearing intracranial luciferase-expressing CTD tumors were randomized 1 week after implantation to receive vehicle (p.o.), anti-PD-1 alone (10 mg/kg, i.p.), or combination therapy consisting of regorafenib (10 mg/kg, p.o.) and mirdametinib (0.5 mg/kg, p.o.), with or without anti-PD-1 (10 mg/kg, i.p.). Regorafenib and mirdametinib were administered three times per week, whereas anti-PD-1 antibody was administered once weekly. Treatment continued for 6 weeks. Tumor progression was monitored weekly by IVIS imaging beginning 1 week after implantation. **j** Kaplan–Meier survival curves for mice treated as described in **i**. Median survival times (days) were 18.5, 26, 27, and 62.5 for the vehicle, anti-PD-1, MIR_REG, and MIR_REG_PD1 groups, respectively. Group sizes (n) are indicated in the risk table; log-rank test results: **P < 0.01, ***P < 0.001, ns, not significant. **k** CellChat-inferred ligand–receptor communication probabilities from monocytes to Teffector and Treg cells.

Next, we performed clustering analysis of published MB patient scRNA-seq data^67, 68, 71^ (**Supplementary Table 9**), and then used our developmental reference to assign developmental stages to each cluster. To do so, we extracted the malignant MB cells from the GSE155446 dataset (10,537 cells from one WNT, nine SHH, and 18 G3/G4 MB patients; **Supplementary Fig. 7c and 7d**; **Supplementary Table 10**), annotating the clusters using marker genes for the seven developmental stages of the glutamatergic lineage (**Fig. 5c and Supplementary Fig. 7e**). Our mapped results revealed a predominance of GCP-like cells in SHH tumors, UBC_pro and UBC_GCP_pro cells in G3 tumors, UBC-like cells in G4 tumors, and RL_pro-like cells in WNT tumors. Notably, the UBC_GCP_pro cluster closely resembled the NSC-like cells reported in the original study (**Supplementary Fig. 7d and 7e**). These findings, which support distinct developmental origins for the major MB subtypes (**Fig. 5d**), were later validated using two independent scRNA-seq datasets^67, 71^ (**Supplementary Fig. 7f-i**; **Supplementary Table 11-12**). The analyzed GSE155446 data, including gene expression and activity profiles, are available through an interactive portal (https://scminer.stjude.org/study/GSE155446).

In addition to the scRNA-seq analysis, we also employed our recently developed ReDeconv^77^ algorithm to deconvolute bulk RNA-seq data from MB patient cohorts and PDX models (see **Methods**). Analysis of the mean percentage of each developmental state among patients with the same tumor subtype^9, 63, 78^ revealed the same patterns observed in the scRNA-seq data (**Supplementary Fig. 8a**): SHH tumors were enriched with GCP-like cells, G3 tumors with UBC_pro and UBC_GCP_pro cells, and G4 tumors with UBC-like cells.

Next, we investigated the activity of BRAF and MAPK11, the primary targets of MEK inhibitors and regorafenib in G3 MB, across lineage stages. Using our scMINER algorithm^79^ and the Mbi, we calculated driver activity at the single-cell level and found markedly elevated combined BRAF-MAPK11 activity in the UBC_GCP_pro and UBC_pro-like subpopulations that are predominant in G3 tumors. No such elevation was observed for G3 minority subpopulations like GC and UBC, which represent more differentiated developmental states (**Fig. 5e**). The drug signature scores for regorafenib + MEK inhibitor showed a similar pattern of elevation in UBC_GCP_pro and UBC_pro-like cells. In contrast, both BRAF and MAPK11 activity and the drug signature scores were minimal in those patients’ normal immune cells. Notably, expression of BRAF and MAPK11 was low across all malignant cell types, demonstrating the usefulness of the activity-based approach (**Supplementary Fig. 8b-d**). As expected, the drug signature scores were low in normal human cerebellum. Although BRAF and MAPK11 expression remained low in normal cerebellar cells, combined BRAF+MAPK11 *activity* was markedly elevated in the UBC_GCP_pro population (**Supplementary Fig. 8e and 8f**).

These findings led us to hypothesize that the regorafenib-MEK inhibitor combinationtargets the putative developmental-origin populations of G3 MB, namely UBC_GCP_pro and UBC_pro cells. To test this hypothesis, we performed scRNA-seq profiling of D341 cells both prior to implantation in mice and then after implantation and treatment with vehicle control, mirdametinib, regorafenib, or their combination. Analysis of the pre-implantation D341 cells revealed high proportions of UBC_GCP_pro and UBC_pro-like subpopulations, much like the G3 MB patient samples (**Supplementary Fig. 8g and 8h**). D341 lacked mature-like populations (GC and UBC) and exhibited slightly higher proportions of GCP-like cells, likely due to culture conditions favoring proliferative progenitor-like states. The combined driver activity and drug signature scores were again exceptionally high in the UBC_GCP_pro-like tumor cells (**Supplementary Fig. 8i**).

Consistent with the *in vitro* findings, D341 tumor cells isolated from treated mice consisted predominantly of UBC_GCP_pro- and UBC_pro-like cells (**Fig. 5f and 5g**; **Supplementary Table 13**). Driver activity was reduced in tumors from the MIR and MIR+REG treatment groups, and drug signature scores were significantly lower in the MIR+REG group than in either the vehicle control or single-agent treatment groups (**Fig. 5h and 5i**). Importantly, treatment effects were cell-state specific. The UBC_GCP_pro-like population demonstrated higher baseline driver activity and greater treatment sensitivity, resulting in a reduced UBC_GCP_pro-to-UBC_pro cell ratio in the MIR and MIR+REG groups (**Fig. 5g** and **5j**). Notably, the emergence of NSC-like tumor cells following combination treatment may represent a potential mechanism of therapeutic resistance.

scRNA-seq analysis of tumors from the SJMB016880 preclinical study (**Fig. 4c**) yielded results consistent with those observed in the D341 model. Regardless of treatment condition, all samples were dominated by UBC_GCP_pro- and UBC_pro-like cell populations (**Supplementary Fig. 9a** and **9b**). Both BRAF+MAPK11 activity and drug signature scores were significantly reduced in the combination-treatment group compared with the vehicle-control and single-agent treatment groups (**Supplementary Fig. 9c** and **9d**). Consistent with the D341 data, the dominant UBC_GCP_pro- and UBC_pro-like populations exhibited higher baseline driver activity and greater drug-sensitivity scores than other cell states (**Supplementary Fig. 9e**).

Compared with the D341 model, the SJMB016880 model contained a larger proportion of more differentiated cell populations, including a substantial UBC-like population and a smaller GC-like population (**Supplementary Fig. 9e**). Although the combination-treatment group exhibited a higher proportion of NSC-like cells after normalization to the total number of cells in each sample (**Supplementary Fig. 9f**), the overall NSC-like population remained markedly lower than that observed in the D341 model. In addition, a distinct RL_pro-like cell population was identified in the SJMB016880 model, and its proportion was enriched in the MIR+REG treatment group. These findings suggest that both NSC-like and RL_pro-like states may contribute to adaptive responses following combination therapy.

Our single-cell analyses traced the major cellular origins of G3 MB to UBC_GCP_pro and UBC_pro cells within the glutamatergic lineage of developing cerebellum. Furthermore, *in vivo* single-cell profiling demonstrated that these populations exhibit elevated driver activity and drug signature scores relative to other G3 MB tumor-cell states, and that combination treatment significantly suppresses both driver activity and drug signature scores across tumor cells. At the same time, combination treatment was associated with the emergence of NSC-like and RL_pro-like populations with potential treatment resistance, highlighting possible cellular mechanisms underlying residual disease and therapeutic escape.

### Regorafenib plus MEK inhibitor is effective and reprograms the suppressive TME in an immunocompetent syngeneic mouse model of G3 MB

While our *in vivo* studies of human cell line xenografts (D341) and PDX models (SJMB016880) were effective in establishing the efficacy of regorafenib-MEK inhibitor combinations in G3 MB tumors, these models require immune-deficient CD-1 nude mice, making them unsuitable for investigating drug treatment in the context of the tumor immune microenvironment (TME). To address this limitation, we utilized CTD, a syngeneic G3 MB mouse model^74^, to evaluate both the therapeutic efficacy of the combination in an immunocompetent setting and its effects on the TME.

CTD is a genetically engineered mouse model that resembles human G3 MB both histologically and molecularly^80^. In the CTD model, *Ctdnep1,* one of the most frequently altered tumor suppressors in G3 MB, is conditionally knocked out in neuronal progenitor cells or cerebellar stem cells, leading to MYC amplification and p53 inactivation, both hallmarks of G3 MB^80^. To confirm the faithfulness of the model, we used scRNA-seq to characterize cultured CTD tumor cells. Most of the CTD tumor cells were UBC_GCP_pro and UBC_pro-like, recapitulating the developmental origins of G3 MB (**Supplementary Fig. 10a**). Consistent with our findings in human G3 MB samples, the UBC_GCP_pro and UBC_pro-like cells had higher BRAF and MAPK11 activity and drug signature scores than RL_pro-like cells (**Supplementary Fig. 10b**). These results suggested that the CTD model was suited to our needs.

We next assessed the efficacy and synergy of the drug combination in CTD cells *in vitro*. Similar to the findings in the human SJMB016880 model (**Fig. 4b**), the CTD cells exhibited marked sensitivity to MEK inhibitors, with low doses of MEK inhibitor significantly reducing the EC_50_ of regorafenib (**Fig. 6a and 6b**). Because CTD cells were more sensitive to MEK inhibition, the concentration range of mirdametinib was reduced by 10-fold for synergy testing. Under these conditions, the combination achieved a mean ZIP score of 12.66, indicating significant synergistic effect (**Supplementary Fig. 10c**). Consistent with observations in human G3 MB cell lines (**Fig. 2e**), combination therapy simultaneously downregulated phosphorylated MEK (p-MEK), ERK (p-ERK), p38(p-p38), and BRAF (p-BRAF), while total protein levels remained unchanged (**Fig. 6c**).

We then evaluated the therapeutic efficacy of this combination *in vivo* using CTD-bearing C57BL/6 mice (**Fig. 6d**). Mirdametinib was administered at a low dose (0.5 mg/kg) because of the pronounced sensitivity of CTD cells to MEK inhibition *in vitro* and its favorable brain penetration. Tumor volume was monitored via MRI weekly, and the relative tumor burden was assessed four weeks post-surgery. Among all treatment groups, only the combination therapy produced a significant reduction in tumor burden relative to vehicle-treated controls from day 14 onward (**Fig. 6e** and **Supplementary 10.d**). Likewise, combination therapy significantly prolonged overall survival compared with both monotherapy and control groups (**Fig. 6f**).

To investigate the effect of combination therapy on tumor cell states and the immune microenvironment, we performed scRNA-seq of endpoint tumor samples from each therapy group. To enhance the representation of TME populations, CD45-based cell sorting was used to enrich immune cells prior to sequencing, targeting an approximately equal proportion of CD45^+^ and CD45^-^ cells (**Fig. 7a**). A total of 39,658 cells were profiled, encompassing both tumor cells and multiple TME populations, including macrophages, monocytes, microglia, dendritic cells, T cells, natural killer cells, neutrophils, basophils, endothelial cells, and oligodendrocyte progenitor cells (**Fig. 7b and Supplementary Fig. 10e**)

CTD tumors treated with regorafenib, both alone and in combination with mirdametinib, had a lower proportion of UBC_GCP_pro-like and UBC_pro-like tumor cells and a higher proportion of RL-pro-like cells compared to control and mirdametinib-treated tumors (**Fig. 7c and 7d**). Consistent with these changes in cellular composition, tumor cells from the REG and MIR+RGE groups showed reduced overall driver activity and drug signature scores (**Supplementary Fig. 10f**). Not surprisingly, RL-pro-like tumor cells exhibited lower driver activity and drug signature scores than the drug-sensitive UBC_GCP_pro and UBC_pro-like populations (**Supplementary Fig. 10g**). Altogether, these findings suggest that REG or MIR+REG treatments selectively reduce drug-sensitive UBC_GCP_pro-like and UBC_pro-like populations while enriching more drug-tolerant RL-pro-like cells.

To investigate whether drug treatment induced additional transcriptional changes in tumor cells, we performed differential expressed gene (DEG) analysis on the scRNA-seq data. Combination-treated tumor cells displayed increased expression of major histocompatibility complex (MHC) genes (*H2-D1*, *H2-K1, H2-Aa, H2-Ab1, B2m* and *H2-Eb1*) together with reduced expression of *Lgals3*, which encodes a ligand for the immune checkpoint receptor LAG3 expression on T cells^81^ (**Fig. 7e**). Functional enrichment analysis of upregulated genes (avg_log2FC > 1, adjusted p value < 0.05) revealed significant enrichment of pathways associated with antigen presentation^82^ (**Fig. 7f**). We generated an antigen-presentation module score and found increased scores across treatment groups, with the highest values occurring in the combination-treatment group (**Supplementary Fig. 10h and 10i**).

Next, we compared immune-cell compositions across treatment groups using the scRNA-seq data (**Supplementary Table 14**). Sixteen immune cell types were captured, including seven T cell populations (**Fig. 7b and Supplementary Fig. 10j**). Compared with control and monotherapy groups, combination therapy increased the abundance of Monocytes (Mono) and T effector cells (Teffector) while reducing T regulatory cells (Treg) (**Fig. 7g**). Notably, the combination-treatment group also displayed increased macrophage infiltration; however, a larger fraction of macrophages exhibited an M1-like phenotype (**Supplementary Fig. 11a**; **Supplementary Table 14**), consistent with a previous report showing that regorafenib enhances M1/M2 polarization in head and neck cancers^83^. In terms of T cells, we found a reduction in immunosuppressive Tregs and an increase in CD8+ T effector cells in the combination therapy group; in fact, this group had the highest ratio of CD8+ T effector cells to Tregs (**Fig. 7h**), indicating more robust anti-tumor immunity compared to the other groups. These changes are consistent with a more immunostimulatory TME and enhanced antitumor immune activity.

To assess the direct impact of therapy on immune cells, we calculated drug signature scores for immune populations. Immune cells from the combination therapy group exhibited the highest regorafenib and mirdametinib signature scores, indicating substantial transcriptional remodeling of the TME by treatment (**Supplementary Fig. 11b**, see **Methods**). Tregs had the highest scores for both drugs; T effector cells and monocytes also presented high scores (**Supplementary Fig. 11c**). These observations suggest that the combination therapy may directly influence both the composition and functional state of immune cells within the tumor microenvironment.

We also found that CD8+ T cells (Tex, T effector cells, Tcm and proliferative T cells) were more activated, as indicated by increased expression of *Ifng*, *Prf1*, in the combination treatment group (**Supplementary Fig. 11d**). These T cells also expressed more *Ccl3* and *Ccl4*, chemokines associated with T cell activation^84, 85^. CCL3 and CCL4 secretion has also been reported in T effector cells activated by infections^86^. In addition, CD8+ T cells from the combination therapy group demonstrated increased expression of *Cxcr6* and reduced expression of *Cxcr4*, a transcriptional profile linked to improved T-cell recruitment and retention within tumors^87, 88, 89,90^. Together, these findings indicate that combination therapy is associated with robust CD8+ T cell activation and antitumor immunity.

Interestingly, CD8+ T cells in the combination treatment group also showed increased expression of several immune checkpoints and immunotherapy targets^91, 92^, including PD1, LAG3, TIM3, CD160, and CD39, as well as an elevated composite immunotherapy-target gene module score (**Supplementary Fig. 11e and 11f**). These findings suggest the emergence of an exhausted T-cell phenotype, which may contribute to residual disease and therapeutic resistance. Collectively, these observations raise the possibility that combining immune checkpoint blockade with regorafenib–MEK inhibitor therapy could further enhance antitumor efficacy.

To test this hypothesis, we conducted an additional preclinical study incorporating anti-PD-1 therapy into the mirdametinib–regorafenib (MIR+REG) treatment regimen. Because the parental CTD model exhibits an engraftment efficiency of approximately 60%–80%, we improved model consistency by isolating luciferase-expressing CTD cells, serially passaging them in immunocompetent mice, and establishing a CTD-luciferase model with 100% engraftment efficiency. Notably, the triple-combination regimen significantly prolonged survival compared with both the MIR+REG dual-combination therapy and the control groups (**Fig. 7i and 7j**). We also observed that mice in the vehicle-treated CTD-luciferase cohort had a shorter median survival than mice bearing the original CTD (luciferase-negative) tumors (18.5 versus 35 days, respectively), suggesting that the CTD-luciferase model may represent a more aggressive disease phenotype than the parental CTD model.

Previous studies have shown that expression of CCL3 and CCL4 in T cells can also be induced by IL12, IL18 and IFNg^86, 93^. Among TME populations, monocytes exhibited the highest expression of *Il18*, while its receptors (*Il18r* and *Il18rap*) were highly expressed on T cells (**Supplementary Fig. 11g-j**). Given the increased abundance of monocytes in the combination treatment group, enhanced monocyte-derived IL18 signaling may contribute to T cell activation by promoting the expression of CCL3 and CCL4 in CD8+ T cells (**Supplementary Fig. 11k**). This hypothesis was partially supported by CellChat analysis^94^. Effector T cells in the combination treatment group received increased IL18-mediated signaling interactions, whereas no comparable increase was observed in Tregs (**Fig. 7k**, see **Methods**). For the CCL3/4–CCR5 axis, signaling toward effector T cells increased in both monotherapy and combination treatment groups, with the strongest interactions observed in the latter. Notably, MIR treatment blocked CCL3/4-CCR5 communications received by Tregs (**Fig. 7k and Supplementary Fig. 11k**). Collectively, these findings support a model in which combination therapy selectively enhances IL18- and CCL3/4-dependent signaling in effector T cells while limiting immunosuppressive Treg activity, thereby suggesting potential engagement of antitumor immunity.

Taken together, regorafenib and mirdametinib combination therapy demonstrated robust antitumor activity in an immunocompetent syngeneic model of G3 MB while simultaneously remodeling the tumor immune microenvironment. The combination therapy was associated with CD8+ T cell activation and antitumor immune responses, whereas the emergence of checkpoint-expressing exhausted T cells suggested a potential mechanism of adaptive resistance. The additional survival benefit observed with anti-PD1 treatment further supports the therapeutic rationale for combining regorafenib–MEK inhibition with immune-checkpoint blockade in G3 MB.

Several limitations of this model and analytical approach should be considered. Although the CTD model provides a valuable immunocompetent platform for evaluating therapeutic responses, it does not fully recapitulate the immune heterogeneity, HLA-restricted antigen presentation, or the complex immunosuppressive features of the human tumor microenvironment^95, 96^. In addition, previous studies using humanized mouse models have reported that the immunomodulatory effects of regorafenib may differ between mice and humans^97, 98^. Potential differences in the tumor microenvironment between the cerebellum and cerebral cortex^69, 99^ should also be considered when interpreting these findings. Accordingly, the immune-related findings reported here warrant further evaluation in humanized preclinical models and clinical specimens. Furthermore, the single-cell transcriptomic and CellChat analyses are primarily descriptive and infer cell-state transitions and intercellular communication patterns computationally. Consequently, they do not provide direct evidence of functional immune activation. Validation of these observations will require orthogonal approaches, including flow cytometric characterization, cytokine profiling, CD8⁺ T-cell depletion experiments, and spatially resolved imaging analyses.

## DISCUSSION

While systems biology has previously been used to identify potential drug candidates in medulloblastoma^100^, to our knowledge this study is the first to apply a systems biology platform to prioritize combination drug screening candidates specifically for high-risk medulloblastoma. The SINBA platform integrates gene activity inference, a brain-penetrating drug database, and single-agent high-throughput screening results to nominate synergistic drug pairs for combination screening. Through single-agent screening, we identified both novel and previously reported G3 MB-specific drugs, such as mycophenolate mofetil and digoxin^100^. Applied to transcriptomic data from three MB patient cohorts, SINBA prioritized a regorafenib-MEK inhibitor combination for G3 MB; these drugs were later experimentally validated to be synergistic both *in vitro* (through drug screening) and *in vivo* (through efficacy studies). We found that tumor cells sensitive to the regorafenib-MEK inhibitor combinations share transcriptomic signatures with early unipolar brush cell (UBC) progenitors, which have previously been proposed as the cell-of-origin for G3 MB^67, 69, 70, 101, 102^. We also showed that a regorafenib-MEK inhibitor combination reprograms the tumor microenvironment and is associated with CD8+ T cell recruitment in an immunocompetent G3 MB mouse model.

G3 medulloblastoma is an aggressive cerebellar tumor arising from malignantly transformed early UBC progenitors within the rhombic lip subventricular zone (RL^svz^)^69, 70, 102^. Single-cell analyses comparing MB cells to normal cerebellar references have found that MB cells arise from early developmental arrest^103^ driven by genetic mutations or epigenetic dysregulation^69, 9, 104^. Novel immunotherapies, such as CAR-T cells targeting Protogenin-expressing stem cells, have shown efficacy in preclinical G3 MB models in immunocompromised mice^102^. However, drug repurposing strategies that target origin cell populations *in vivo* are lacking. While SINBA is not expressly designed to select synergistic drugs that target originating populations, it may nonetheless propose drug pairs that do so, as it did for G3 MB.

Through the SINBA algorithm, trametinib and regorafenib were prioritized as a candidate synergistic drug combination targeting BRAF and MAPK11 (a p38 subunit), two putative drivers of G3 MB. Although SINBA does not explicitly model the non-linear interactions (e.g., feedback loops, adaptive resistance, lineage-state transitions) that may underlie biological synergy, the synergistic activity of this drug pair was subsequently identified and validated through both *in vitro* and *in vivo* studies. While this combination has not previously been evaluated in MB, synergistic effects between MEK inhibitors and regorafenib have been reported in colorectal cancer^105, 106^, gastric cancer^107^ and melanoma models harboring BRAF V600E mutations^108^. Although BRAF mutations are rare in MB, elevated MEK signaling is frequently observed, even in the absence of changes in total protein expression^50^. Importantly, mirdametinib and regorafenib converge on the MAPK signaling pathway, which has been implicated in medulloblastoma progression and metastasis. Activation of the PDGFA/RAS/MAPK signaling pathway has been linked to MB metastasis^109, 110^, and pharmacological inhibition of MEK1/2 with U0126 reduces tumor cell viability and metastatic potential ^104,111^. Additional evidence supports the relevance of MAPK signaling in treatment-resistant MB. Recurrent MB cells maintain a stem cell-like state through overexpression of the BPIFB4 (bactericidal/permeability-increasing fold-containing family B member 4), and BPIFB4 knockdown reduces tumor aggressiveness *in vitro* and *in vivo*. Notably, ERK1/2 and p38α were among the proteins most affected by BPIFB4 depletion^112^. Consistent with these findings, inhibition of ERK and p38-MAPK signaling by liposomal honokiol induces reactive oxygen species (ROS)-mediated apoptosis in MB cell line models^113^. Together, these studies provide a strong biological rationale for dual targeting of MEK and p38-MAPK signaling in G3 MB. Beyond suppressing tumor growth, the mirdametinib–regorafenib combination may also have the potential to limit metastatic progression, a possibility that warrants further investigation in preclinical metastasis models.

Robust clinical and preclinical evidence suggests that combining regorafenib with MEK inhibitors may be effective for the treatment of brain tumors^114, 115, 116, 117^. In the phase II REGOMA trial, regorafenib demonstrated superior therapeutic outcomes compared with lomustine in patients with recurrent glioblastoma^28^. The MEK inhibitor trametinib, in combination with the BRAF inhibitor dabrafenib, received FDA approval in 2022 for the treatment of advanced tumors harboring BRAF V600E mutations, including adult and pediatric gliomas. Another MEK inhibitor, mirdametinib, has recently received FDA approval for the treatment of plexiform neurofibromas in both adult and pediatric patients. Notably, the combination of mirdametinib and the PI3K/mTOR inhibitor paxalisib significantly extends survival in mouse models of pediatric high-grade gliomas^58^. Given its more favorable pharmacokinetic profile and CNS activity, mirdametinib was selected as the MEK inhibitor for our *in vivo* studies.

The regorafenib-MEK inhibitor combination inhibits cancer progression both directly, by targeting cancer-driving genes, and indirectly, through modulation of the TME. MEK inhibition protects tumor-infiltrating CD8+ T cells from exhaustion driven by chronic TCR stimulation while preserving cytotoxic activity^118^. Regorafenib promotes a tumor-suppressive microenvironment by enhancing M1 macrophage polarization, limiting M2 polarization, reducing PD-L1 expression, and increasing MHC class I gene expression in tumor cells^83, 97, 119^. These observations are consistent with our finding that regorafenib-MEK inhibitor therapy is associated with an antitumor TME in G3 MB. The upregulation of immune checkpoint molecules on CD8+ T cells following combination therapy may contribute to treatment resistance, while also providing a potential therapeutic opportunity through the addition of immune checkpoint blockade, such as anti-PD-1/PD-L1, LAG3, or TIM3-targeted therapies. Consistent with this hypothesis, we found that adding anti-PD1 therapy to the combination regimen further significantly prolonged survival in the syngeneic mouse model. At present, no immunotherapy has been approved for G3 MB tumors, which is generally considered immunologically “cold” and largely resistant to immunotherapeutic approaches. By promoting a more immunostimulatory tumor microenvironment, regorafenib-MEK inhibitor therapy may sensitize G3 MB tumors to immune checkpoint blockade, providing a potential strategy to extend the benefits of immunotherapy to this patient population.

The primary goal of this study was to identify and prioritize novel synergistic drug combinations with strong clinical translational potential for high-risk MB, rather than to formally benchmark the predictive performance of the SINBA framework. Using this strategy, we identified and validated the therapeutic efficacy of the regorafenib–mirdametinib combination in both *in vitro* and *in vivo* models. Importantly, these findings should be interpreted as evidence supporting the efficacy of a prioritized drug combination rather than a definitive validation of the predictive performance of SINBA itself. SINBA does not explicitly model non-linear driver–driver interactions, such as cooperation, antagonism, or synthetic lethality. Future development of the framework could incorporate interaction-aware features, including driver interaction terms and perturbation-based modeling strategies, to better capture complex biological dependencies and further refine candidate prioritization. A rigorous evaluation of SINBA’s predictive performance would require prospective, blinded testing of multiple high- and low-ranked drug combinations across molecularly diverse preclinical models to assess robustness and generalizability, including MYC-high and MYC-low tumors, as well as treatment-naïve and relapsed disease models. Given the substantial molecular and clinical heterogeneity of MB, biomarker-guided patient stratification will likely be essential for future clinical validation efforts. While these studies represent important future directions, they fall outside the scope of the present work, which was designed as a therapeutic discovery study rather than an algorithm-validation study. In addition, although we focused on prioritized small-molecule drug pairs with favorable BBB penetration, ongoing advances in drug delivery technologies^120^ may expand the range of therapeutically actionable compounds and combinations that can be evaluated using this framework.

## MEHTODS

### Chemicals

Regorafenib monohydrate (HY-10331A), Trametinib (HY-10999), and Mirdametinib (PD0325901, HY-10254) were obtained from MedChemExpress (MCE). For in vitro studies, each compound was dissolved in dimethyl sulfoxide (DMSO; FisherScientific, BP231-100) to a final stock concentration of 10 mM. For in vivo preclinical studies, the compounds were formulated as follows. 1. Regorafenib: The free base equivalent of regorafenib was calculated using a correction factor of 1.0373 for its monohydrate form. The drug was first dissolved in an 80% non-aqueous vehicle consisting of propylene glycol (PG, 42.5%), polyethylene glycol 400 (PEG400, 42.5%), and Pluronic F68 (15%) (v/v/v) by heating to 60 °C with agitation and vortexing. The solution was then brought to final volume with ultra-pure water at room temperature, yielding a clear formulation for oral gavage at a concentration of 3 mg/mL, administered at 10 mL/kg. 2. Trametinib: A 3 mg/mL stock solution was prepared in DMSO. For oral administration, the stock was diluted to 1% DMSO in a vehicle consisting of 10% Cremophor RH40, 10% PEG400, and 80% ultra-pure water, resulting in a final dose concentration of 0.03 mg/mL, delivered at 10 mL/kg. 3. Mirdametinib: The compound was formulated at 2.5 mg/mL in an aqueous suspension containing 1% methylcellulose and 1% Tween 80. The formulation was prepared using sonication to ensure uniform dispersion and administered by oral gavage at a dose volume of 10 mL/kg.

### The SINBA algorithm to identify synergistic drug pairs for G3 MB

The SINBA package is available on GitHub, along with comprehensive documentation and a step-by-step tutorial (https://github.com/jyyulab/SINBA). To enable prioritization of synergistic drug combinations with favorable blood–brain barrier (BBB) permeability, we integrated the Brain Penetrant Drug–Target Database (BPdb) into the SINBA framework. BPdb drugs were classified into three categories: (1) experimentally validated BBB-penetrant small molecules with reported CNS exposure; (2) small molecules with putative BBB permeability, supported by their evaluation in clinical trials for neurological disorders, primary brain tumors, or brain metastases; and (3) antibody-based therapeutics considered potentially CNS-accessible based on emerging evidence of intracranial activity and advances in CNS-targeted drug delivery technologies. Collectively, these categories include both established and candidate therapeutic agents with potential applicability to brain tumors. Below, we describe the workflow used to identify synergistic therapeutic candidates for Group 3 medulloblastoma (G3 MB) using SINBA, focusing on Category 1 and Category 2 small molecules. Because BBB permeability has not been experimentally confirmed for all Category 2 compounds, additional validation will be required to establish their CNS penetration properties.

#### 1. Construction of medulloblastoma-specific gene-gene interaction network

To reconstruct medulloblastoma-specific interactomes (MBi), we applied the SJARACNe algorithm^44, 121^ to a curated microarray dataset, DKFZ_SJ, comprising 407 medulloblastoma patient samples. Using Gene Ontology classifications, we compiled a list of 11,239 transcription factor (TF) genes and signaling molecules to serve as hub genes in the MBi. Separate transcription factor (TF) and signaling molecule (SIG) networks were generated using SJARACNe. In this final network, drivers were linked to their downstream targets through edges based on gene–gene mutual information derived from their expression patterns. The reconstructed MBi consisted of 38,621 probe nodes and 1,567,381 interactions, representing 12,495 hub probes (corresponding to 7,155 genes) and their downstream target probes.

#### 2. Identification of G3-Specific Drivers

Hub gene activities for each sample in the DKFZ_SJ dataset (407 MB patients, 4 normal cerebellum) were inferred using NetBID2 package^48^ function **cal.Activity** (es.method = “weightedmean”), which integrates gene expression profiles with the MBi. G3-specific drivers were selected based on the following criteria, resulting in 75 candidates:

- Adjusted p-value for G3 vs. other subgroups < 1e-5, with Z-score > 0.
- Adjusted p-value for G3 vs. normal cerebellum < 1e-3, with Z-score > 0.
- Regulon size between 50 and 500.

#### 3. Refinement of G3-Specific Drivers

To refine the list, we used SINBA’s built-in Brain Penetrant database (BPdb) to filter for druggable drivers, yielding 45 seed drivers targeted by 69 compounds. Thus, seed drivers were selected based on both inferred biological activity and druggability.

The TF and SIG networks were merged using the SINBA function **get.combined.network** to prepare for activity calculations and driver-pair predictions. Driver pairs were generated either manually or by unbiasedly combining seed drivers with all druggable hub genes using the **combineNet2Target_list** function.

Following seed drug library screening, 20 drugs (targeting 20 seed drivers) with EC20 < 10 μM or the solo agent targeting on the interesting seed drivers.

#### 4. Prediction of Driver Pairs and Drug Pairs

Driver-pair predictions were performed by combining the TF and SIG networks and pairing the 20 seed drivers with all druggable MBi hub genes using the **combineNet2Target_list** function. Activities for single drivers and driver pairs were calculated using **cal.SINBA.Activity**. Driver pairs were ranked using **get.SINBA.DE.2G**, which computes statistical values comparing G3 with other subtypes and normalizes p-values into Z scores (Z.pair). Z scores for seed (Z.seed) and partner drivers (Z.partner) were calculated similarly. The delta.Z score was defined as:

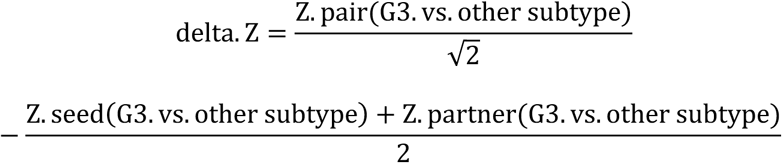

A genetic algorithm^52^ (implemented in the function **find_max_sum_submatrix**) was then employed to optimize the maximum sum of delta.Z scores (*in silicon* synergistic scores) for G3 vs. other subtypes. The final matrix layout consisted of 20 seed drivers × 16 partner drivers. BPdb was then used to convert driver pairs into drug pairs.

### SINBA-guided drug combination screening

#### 1. Single-Drug Dose–Response Testing

A total of 90 compounds (69 predicted seed drugs and 21 partner drugs) identified by the SINBA algorithm were screened. Compounds were tested in 10-point dose-response curves (10 µM to 0.47 nM, 1:3 serial dilution) against the HDMB03 cell line. Cell culture conditions, tumor dissociation protocols, and density optimization followed established methods^21^. HDMB03 cells were seeded in 384-well plates. After 24 hours, compounds were pin-transferred (Biomek FX Workstation; Beckman Coulter) with negative (0.08% DMSO) and positive control (1.6 µM staurosporine; MedChemExpress, HY-15141). Cell viability was assessed 7 days post-treatment using CellTiter-Glo® (Promega, G7572). Dose-response curves were fitted using the drc package^122^ with four-parameter log-logistic function to estimate EC_20_, EC_50_ values, which defined low- and high-dose conditions (with fixed defaults of 1 μM and 6 μM, respectively, for drugs lacking EC_20_ or EC_50_ values).The four-parameter log-logistic function is given by the expression:

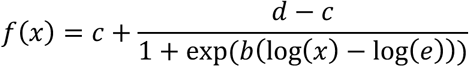

#### 2. Two-dose combination testing

To prioritize the drug pairs testing for synergy, 320 SINBA-predicted drug pairs were tested in 384-well plates (64 wells reserved for controls). Each pair included four conditions: drug1 low dose+ drug2 high dose; drug1 high dose + drug2 high dose; drug1 low dose + drug2 low dose; drug1 high dose + drug2 low dose. 7 days after treatment, viability was measured (CellTiter-Glo®, Promega) and normalized to controls. Combination effects (Combo_effect_ or Δinhibition_mean_) were quantified as:

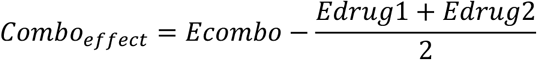

Δinhibition_max_ measures the difference in inhibition between a drug combination and the more effective of its two constituent drugs when administered as single agents:

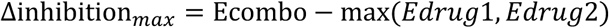

To reduce the likelihood of false-positive identification of synergistic drug pairs during subsequent screening, we excluded drug pairs that achieved >95% maximal growth inhibition while exhibiting a Δinhibition_max_ <10%. Such combinations are often driven by the strong activity of the individual agents and therefore tend to reflect additive, rather than truly synergistic effects.

#### 3. 10-dose combination screening

32 prioritized pairs were tested in full dose-response matrices (10×10 concentrations, 10 umol/L to 0.14 nmol/L). Drugs were added simultaneously, and viability was assessed after 7 days. Synergy score were calculated using the synergyFinder package^123^ with zero interaction potency (ZIP) model^35^.

Synergy was assessed using a standardized dosing scheme applied across all drug combinations. This approach was selected to accommodate the practical constraints of our in-house high-throughput screening platform while maintaining a focus on combinations with potential clinical relevance. However, tailoring dose ranges to the pharmacologic properties of individual agents, such as their EC₅₀ values, may improve the sensitivity of synergy detection and increase the identification of promising drug pairs. Accordingly, we recommend that future applications of SINBA incorporate either (i) an adaptive dose design based on each drug’s measured EC₅₀, or (ii) an initial broad-range dose matrix for novel or uncharacterized drug pairs.

### Medulloblastoma Patient SINBA Shiny App

We developed a medulloblastoma-specific SINBA Shiny app to provide a comprehensive resource for exploring MB patient data(https://yulab-stjude.shinyapps.io/SINBA_MB). The app includes pre-built MBi networks, curated six drug-gene interaction databases, driver and driver-pair activity profiles, gene expression profiles, and patient clinical outcomes. Users can flexibly select druggable drivers using six integrated drug-gene interaction databases and explore potential synergistic driver pairs. This tool is designed to support hypothesis generation and facilitate the identification of therapeutic targets in medulloblastoma research.

### Cell culture and transfection

G3 MB human cell line HDMB03 and MB002 were generously provided by Dr. Till Milde and Dr. Jae-Youn Cho, respectively^124, 125^. PDX model SJMB016880 was utilized as described in the published work^63^. Human SHH subgroup DAOY cell line was purchased from American Tissue Culture Collection (ATCC, HTB-186). Human G3 cell lines D341, D283, D425 and mouse model CTD^80^ were generously provided by Dr. Richard Q. Lu. Human cell lines and PDX model were cultured as neurospheres in supplemented neurobasal medium per established protocol^126^. All lines underwent quarterly mycoplasma testing (MycoAlert™, Lonza) and maintained <10 passages post-thaw.

To generate the luciferase-expressing D341 cell line, cells were infected with a lentiviral vector (CL20SF2-Luc2aYFP) containing the firefly luciferase and yellow fluorescent protein (YFP) genes at a multiplicity of infection (MOI) of 20. Following infection, cells were sorted based on YFP fluorescence to select for positively transduced cells.

The mouse model CTD was cultured using previously published methods. Briefly, CTD cells were grown in DMEM/F-12 medium supplemented with HEPES buffer, sodium pyruvate, non-essential amino acids, GlutaMax (Thermofisher Cat# 35050061) and B-27 supplement (Thermofisher Cat# 17504044) at 37°C and 5% CO2. A total of 1x10^6^ CTD cells were infected with 1µl of virus (titer 4x10^10^) for 48h. Then YFP+ cells were sorted by fluorescent-activated cell sorting (FACS).

### Drug signature score calculation in medulloblastoma tumors

Gene signatures induced by regorafenib and trametinib treatment in the central nervous system were obtained from the iLINCS database (https://www.ilincs.org/ilincs/;regorafenib: CTRS_2230; trametinib: CTRS_2295; Table S15). Differentially expressed genes were identified using cutoffs of |logFC| > 1.5 and p-value < 0.01. This analysis identified 103 upregulated and 42 downregulated genes following regorafenib treatment, and 32 upregulated and 28 downregulated genes following trametinib treatment. The upregulated and downregulated gene sets from both treatments were merged to represent the combined effects of the drug pair.

To calculate MEK inhibitor plus regorafenib signature scores in MB tumor cells, gene expression profiles were first converted to activity profiles by integrating the medulloblastoma-specific interactome (MBi) for bulk or single-cell RNA-seq samples. Activity profiles were standardized using z-transformation. Upregulated and downregulated gene sets were analyzed separately using the NetBID2 function **cal.Activity.GS** (es.method = “ssgsea”). For each sample, the drug pair signature score was calculated by subtracting the ssGSEA score of the upregulated gene set from that of the downregulated gene set, quantifying sensitivity to the combination treatment.

### *Ex vivo* evaluation of MEK inhibitor and regorafenib combination efficacy by AlamarBlue assay

The viability of D341, MB002, D283, D425, DAOY, SJMB016880, and CTD cells were assessed in the presence of MEK inhibitors (trametinib or mirdametinib/PD901) in combination with regorafenib using full dose-response matrices (10×10 concentrations, ranging from 10 umol/L to 1 nmol/L Drugs were transferred into 384-well plates using the Echo 650 series acoustic liquid handler for precise nanoliter-scale dispensing.

Cells were seeded into the drug-treated plates at a density of 1 × 10³ cells per 30 μL of medium and incubated for seven days under standard culture conditions. After seven days, cell viability was measured using the AlamarBlue assay. Briefly, 10% AlamarBlue reagent was added to the cells in complete medium, and the plates were incubated for 1-4 hours according to the manufacturer’s protocol. Absorbance was measured using a plate reader to quantify cell viability. The calculation of synergistic effects was performed as described in the SINBA-guided high-throughput screening section.

### Drug treatment and western blot

Human G3 cell lines or mouse mode CTD cells were seeded into 6-well plates and treated with regorafenib, either alone or in combination with trametinib or mirdametinib, at the indicated concentrations. After 3 days of treatment (72 h), cells were harvested and lysed in RIPA buffer supplemented with protease and phosphatase inhibitors (#78440, Thermo Fisher Scientific) for 20 minutes. Total protein extracts were separated by SDS-PAGE and transferred onto polyvinylidene fluoride (PVDF) membranes (Millipore, USA) using the Trans-Blot Turbo™ transfer system (Bio-Rad, USA).

The PVDF membranes were blocked with 5% non-fat dry milk in TBS-T for 1 h at room temperature. Following blocking, the membranes were incubated overnight at 4°C with primary antibodies (Cell Signaling Technology). After washing three times with TBS-T, the membranes were incubated with HRP-conjugated secondary antibodies (ABclonal) for 1 h at room temperature. Protein bands were visualized using an enhanced chemiluminescence (ECL) kit (ABclonal) and imaged with the Bio-Rad ChemiDoc MP Imaging System.

### Pharmacokinetics of regorafenib, mirdametinib and trametinib in mice

The plasma and brain pharmacokinetic (PK) profile of regorafenib was evaluated in normal female Crl:CD1 (ICR) nude mice (7 to 11 weeks old; Charles River Laboratories, Stock Number 086). Regorafenib monohydrate (MedChem Express, CAT# HY-10331A, LOT# 09764) was suspended in 80% (propylene glycol 42.5% / polyethylene glycol 400 42.5% / Pluronic F68 15%) and 20% ultra-pure water (Millipore), at 1 mg/mL for a 10 mg/kg free base equivalent dose. Terminal samples, one from each mouse, were collected over a 24-hour post-dose period by cardiac puncture using a 1 mL syringe, the blood placed in a Sarstedt Microvette K3EDTA 500 μL tube and immediately separated to plasma. The carcass was then perfused with PBS, the brain extracted, rinsed, and placed in a microcentrifuge tube. All samples were immediately stored on dry ice and transferred to -80 °C until analysis. Remaining regorafenib dosing suspension was submitted for verification of potency, and chemical and physical stability during the study period.

For brain exposure studies of trametinib and mirdametinib, 10 mmol/L stock solutions were initially prepared in DMSO. These stocks were subsequently diluted to a final concentration of 1 mmol/L using a sequential dilution method with 30% polyethylene glycol 400 (PEG400) and 60% phosphate-buffered saline (PBS). Intravenous dosing was performed at 5 mL/kg. Samples were collected as regorafenib study.

Brain exposure was assessed by determining the unbound brain-to-plasma partition ratio (*K*_p,uu_), a critical parameter in evaluating central nervous system (CNS) drug penetration and distribution. *K*_p,uu_, derived from the total brain-to-plasma partition ratio (*K*_p_), was calculated using Equation (1), as previously described^56^. Unbound drug exposures were estimated based on measured total drug concentrations in plasma and brain, in combination with experimentally determined fractions unbound in plasma (*f*_u,plasma_) and brain tissue (*f*_u,brain_). Plasma and brain concentration-time data for the agents were summarized, with the mean values subjected to standard noncompartmental analysis (NCA) using Phoenix WinNonlin 8.1 (Certara USA, Inc., Princeton, NJ). The apparent plasma-to-brain partition coefficient (Kp) in mice was estimated as the ratio of the area under the concentration-time curve (AUC) in brain to the AUC in plasma.

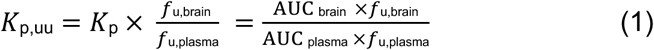

### *In vivo* evaluation the efficacy of mirdametinib plus regorafenib

Luciferase-positive D341 cells (30,000 cells/mouse), SJMB016880 tumor cells (500,000 cells/mouse), DAOY cells (500,000 cells/mouse) and CTD cells (1,000,000 cells/mouse; with or without luciferase expression) were resuspended in 5 µL Matrigel (BD Biosciences, 356234) and stereotactically implanted into the cortices of 7–11-week-old female CD-1 nude mice or male C57BL/6J mice (for the CTD syngeneic model; The Jackson Laboratory, Cat. 000664). Implantations were performed according to a previously established intracranial xenograft protocol routinely used in our laboratory, which has consistently produced reproducible and biologically relevant results ^20, 21, 127^. Previous studies have shown that intracranial medulloblastoma xenografts retain key molecular and pathological characteristics of the original tumors, including histological features, transcriptomic profiles, and epigenetic states, regardless of the specific intracranial implantation site^63^. Nevertheless, potential differences in anatomy (cerebellum versus cerebral cortex) and species-specific features of the tumor microenvironment should be considered when interpreting microenvironment-related findings from these models.

Tumor growth was monitored weekly using bioluminescence imaging (BLI) with a PerkinElmer IVIS imaging system or magnetic resonance imaging (MRI), as previously described^21^. Treatment commenced one week after implantation surgery. Mice were randomized into treatment groups and received continuous treatment for six weeks with vehicle, mirdametinib, regorafenib, or the combination of mirdametinib and regorafenib. Regorafenib was administered at a fixed dose of 10 mg/kg across all models, whereas mirdametinib was dosed at 5 mg/kg in the D341 and DAOY models and 0.5 mg/kg in the PDX SJMB016880 and CTD models. To account for the approximately 80-fold difference in *in vitro* sensitivity to mirdametinib, mice bearing SJMB016880 or CTD tumors received a 10-fold lower dose than D341-bearing mice to achieve comparable pharmacodynamic activity while minimizing potential toxicity. All treatments were administered orally three times per week.

For studies evaluating the combination of regorafenib, mirdametinib, and irradiation in the D341 model, radiation was delivered using the Small Animal Radiation Research Platform (SARRP, Xstrahl Life Sciences). An 8 mmx 11 mm treatment field was delivered to the brain using opposed lateral beams 0.5 Gy/day for 5 days to a total dose of 2.5 Gy. Detailed irradiation protocols have been previously published^128^. A low-dose radiation regimen (0.5 Gy × 5 fractions) was intentionally selected to enrich for radiation-resistant tumor cell populations^129^, while providing a therapeutic window to evaluate whether regorafenib and MEK inhibition could enhance the antitumor efficacy of radiation therapy beyond that achieved by either treatment modality alone.

To investigate potential immune contributions to treatment response, anti-PD-1 antibody was added to the regorafenib–mirdametinib combination regimen in the CTD syngeneic model. Mice received anti-PD-1 monotherapy (10 mg/kg, intraperitoneally) or combination therapy consisting of regorafenib (10 mg/kg, oral gavage), mirdametinib (0.5 mg/kg, oral gavage), and anti-PD-1 antibody. Anti-PD-1 was administered once weekly on the same day as the second weekly dose of regorafenib and mirdametinib.

Mice that reached the experimental endpoint were euthanized, and tumors were harvested and snap-frozen for molecular analysis. Mice were excluded from the study if they died prematurely, failed to develop tumors, or required humane euthanasia due to unrelated health conditions without tumor burden. All experiments were conducted in accordance with the National Institutes of Health’s Guide for the Care and Use of Laboratory Animals and were approved by the St. Jude Children’s Research Hospital (SJCRH) Institutional Animal Care and Use Committee (IACUC protocol #3078).

Toxicity was assessed weekly by measuring body weight and performing endpoint blood work. Complete blood counts were obtained from whole blood treated with ethylenediamine tetra-acetic acid (EDTA) using a Forcyte Hematology Analyzer (Oxford Scientific), which employs impedance and laser technology. Serum diagnostic chemistry panels were analyzed using the Trilogy chemistry analyzer (Drew Scientific).

### Visualization Web Portal for Single-Cell Glutamatergic Lineage Developmental Reference and Medulloblastoma Patient Data

To facilitate data exploration and visualization, we developed two interactive web portals: one for the glutamatergic lineage reference (https://scminer.stjude.org/study/Glutamatergic-lineage) and one for medulloblastoma patient data (https://scminer.stjude.org/study/GSE155446). The portals were built with JavaScript and Vue.js for the frontend, Java and Spring Boot for the backend, and Neo4j and MySQL databases for efficient data management. The web portal provides access to three main functionalities: dataset presentation, data visualization, and data downloading.

For glutamatergic lineage analysis, we curated a public scRNA-seq dataset of the human fetal cerebellum generated with the 10X Chromium platform^71^. From this dataset, glutamatergic lineage cells were filtered by selecting cell populations annotated as “01xNSC,” “06xGCP_progenitor,” “07xGCP_1,” “08xGCP_2,” “09xGranule_cell,” “10xUBC_progenitor,” “11xDiff.UBC,” and “11xUBC” in the original study. To account for major sequencing batch effects, present in the dataset, UMAP-level batch correction was performed using Harmony prior to downstream clustering and visualization. Unsupervised Leiden clustering, combined with previous annotations and manual curation informed by known lineage markers, resulted in the identification of seven distinct glutamatergic lineage cell types: neural stem cells (NSC), rhombic lip progenitors (RL_pro), unipolar brush cell and granule cell precursor common progenitors (UBC_GCP_pro), unipolar brush cell progenitors (UBC_pro), granule cell precursors (GCP), granule cells (GC), and unipolar brush cells (UBC).

For medulloblastoma patient data, single-cell RNA-seq profiles from 28 pediatric tumors (1 WNT, 9 SHH, 7 Group 3, 11 Group 4; GEO accession GSE155446) were processed using the Seurat workflow, with batch correction via Harmony. Cell type annotations were adopted from the original publication or re-annotated using the glutamatergic lineage reference.

Both portals integrate single-cell expression profiles with the medulloblastoma-specific interactome (MBi), utilizing the scMINER algorithm^79^ to generate activity profiles. Users can query both gene expression and activity profiles for any given list of driver genes. The portal supports various visualization tasks, including clustering UMAPs, heatmaps, bubble plots, violin plots, and network plots, enabling users to explore lineage-specific patterns and driver activity in a user-friendly environment.

### Pseudotime trajectory analysis for the glutamatergic lineage cells

The differentiation trajectory of glutamatergic lineage cells was reconstructed using Slingshot^130^ (v2.10) and Monocle 3^131^ (v1.3). The UMAP projection of glutamatergic lineage cells was imported from a Seurat object for downstream analysis. For pseudotime calculation with Slingshot, the slingshot function was applied to compute the arc length from the starting point (neural stem cells, NSC) to all projected points along the principal curve, assigning pseudotime values to each cell. To construct the trajectory with Monocle 3, sample-based batch effects were corrected using the align_cds function. The UMAP projection was imported from the Seurat object, and the differentiation trajectory was generated using the learn_graph function. NSC cells were selected as the starting point for the pseudotime trajectory. The trajectory, aligned with the UMAP projections, was visualized using the plot_cells function in Monocle 3.

### Deconvolution of Glutamatergic Lineage Developmental States in Bulk RNA-seq Samples of MB Patients or PDOX

To prepare the dataset for deconvolution, we normalized the reference glutamatergic lineage single-cell counts to 1 × 10⁶ and used the seven cell type annotations as input for ReDeconv package^77^ built-in functions **get_initial_Signature_Candidates** and **Get_signature_gene_matrix**. The following parameters were applied:

- L_topNo = 200
- L_max_pv = 0.05
- L_min_fold_change = 2.0
- L_CellType_CellNo_LB = 30
- L_NoSep_sampleNo_UB = 2

After generating the glutamatergic lineage cell type signature gene matrix, we applied the **ReDeconv** function to deconvolute glutamatergic lineage developmental state proportions in bulk RNA-seq samples from public medulloblastoma patient datasets and patient-derived orthotopic xenograft (PDOX) bulk RNA-seq data, which were quantified as FPKM. The seven cell type signature matrix served as input for this deconvolution process, allowing us to estimate the relative proportions of developmental states in each bulk RNA-seq sample.

### Single-cell RNA-seq profiling of human cell line D341 and mouse model CTD *in vitro*

Cells from either the mouse syngeneic G3 MB model (CTD) or the human G3 MB cell line (D341) were dissociated with Accutase (Stemcell Technologies, Cat# 07922) at 37°C for 5 minutes. Then we input 8000 cells with 80% viability to the workflow of 3’ gene expression of 10x genomics. The 10x Genomics Chromium controller instrument was used for Gel Bead-in-Emulsion (GEMs) preparation. Gel Bead-in-Emulsion (GEM) generation was performed using the Chromium Controller, followed by single-cell library preparation using Chromium Next GEM Single Cell 3′ Kit v3.1 (PN-1000269), Chromium Next GEM Chip G Single Cell Kit (PN-1000127) and Dual Index Kit TT Set A (PN-1000215). cDNA and final barcoded sequencing libraries were constructed following the manufacturer’s guidelines and subsequently assessed for quality and concentration using Tapestation.

### Single-cell RNA-seq profiling of human model and mouse model after treatment in preclinical study

*In vivo*-implanted D341 mouse tumors were harvested at the endpoint. Tumor tissues were minced and digested at 37°C for 30 minutes using the Human Tumor Dissociation Kit (Miltenyi Biotec, catalog no. 30-095-929). Red blood cells were removed by treatment with ACK lysis buffer (Thermo Fisher Scientific, catalog no. A1049201). The digested tissues were then passed through a 70 μm cell strainer to obtain a single-cell suspension. Approximately 8,000 cells with over 80% viability were selected and processed using the Flex Gene Expression workflow from 10x Genomics for downstream analysis.

Mouse CTD tumors were harvested at end point. Mouse tissues were minced and digested at 37°C for 1 h (1mg/mL papain, 10 μg/mL DNase I, 10% heat-inactivated FBS in RPMI) to dissociate cells. Red blood cells were lysed with ACK lysis buffer (Thermo Fisher Scientific, catalog no. A1049201). Then tissues were passed through 70 μm cell strainer to get single-cell suspension. Then cells were incubated with CD16/32 antibody (Miltenyi Biotec, Cat#130-092-575) for 20 minutes on ice to block Fc receptors. Next, they were stained with PE-CD45 antibody (Biolegend, Cat# 157603) on ice for 20 minutes and DAPI (1ug/ml) on ice for 3 minutes. Then live immune cells (CD45+) and tumor cells (YFP+) were sorted by FACS and combined for downstream library preparations.

Then we input 8000 cells with 80% viability to the workflow of Flex gene expression of 10x genomics. The 10x Genomics Chromium controller instrument was used for Gel Bead-in-Emulsion (GEMs) preparation. Gel Bead-in-Emulsion (GEM) generation was performed using the Chromium Controller, followed by single-cell library preparation using GEM-X Flex Sample Preparation v2 Kit (PN-1000781), GEM-X Flex Gene Expression Chip Kit (PN-1000791), GEM-X Flex Gene Expression Mouse Transcriptome probe kit (PN-1000786) and Dual Index Kit TT Set A (PN-1000215). cDNA and final barcoded sequencing libraries were constructed following the manufacturer’s guidelines and subsequently assessed for quality and concentration using Tapestation.

### Single-cell RNA-seq data analysis

Raw sequencing data generated from the Illumina NovaSeq platform were processed using the Cell Ranger pipeline (version 8.0.1, 10X Genomics). This pipeline performed demultiplexing, alignment to the reference genome (hg GRCh38 or mm39), and barcode processing to produce gene–cell matrices for downstream analysis. To ensure consistent filtering, datasets from in vivo samples treated with either vehicle or drug were merged into a single dataset.

Unsupervised clustering of single-cell RNA sequencing (scRNA-seq) data was performed using the R package Seurat^132^ (version 5.3.0). Cells were filtered based on quality metrics, excluding those with low or high unique molecular identifier (UMI) counts, low or high gene numbers, or elevated mitochondrial gene expression (>10%). Gene expression levels were normalized to 10,000 UMIs per cell and log-transformed by adding 1 to the expression matrix. Highly variable genes (top 2000) were identified using Seurat’s “vst” method, which were subsequently used for principal component analysis (PCA). The first 30 principal components (PCs) were utilized for dimensionality reduction via uniform manifold approximation and projection (UMAP) and clustering. Cell type markers were applied to annotate tumor cell types within each cluster.

### Drug signature score calculation in normal immune cells

Context-matched gene signatures induced by regorafenib and mirdametinib were curated from bone marrow dataset (regorafenib: SCP_87407; mirdametinib: SCP_86814; Table S15) using cutoffs of |logFC| > 1.5 and p-value < 0.05. This yielded 48 upregulated and 55 downregulated genes for mirdametinib, and 58 upregulated and 93 downregulated genes for regorafenib.

For normal cells, gene expression values in each single cell were standardized, and the drug signature score was calculated by subtracting the ssGSEA score of the upregulated gene set from the ssGSEA score of the downregulated gene set for the mirdametinib-regorafenib combination.

### Cell–cell interaction analysis

Cell–cell interaction analysis was conducted using samples from 16 mice (4 mice per treatment arm) profiled with the 10x Flex Gene Expression platform. Interactions were analyzed using CellChat (V2.1.2), focusing on secreted signaling ligand-receptor pairs from the CellChatDB database. The selected cell types included: “Tcm,” “Teffector,” “Tex,” “Tfh,” “Th17,” “Treg,” “Mono,” “Macrophage” “Microglia” and “Tumor” The number and strength of inferred interactions were analyzed between the 10 selected cell types. Differential interaction strength was compared between each treatment arm and the control group. Additionally, the relative interaction strength and information flow for specific receptor-ligand signaling pathways were assessed.

## Supporting information

Supplemental_tabels

## DATA AVAILABILITY

The 10x Genomics single-cell RNA-seq data generated in this study are available from NCBI GEO under accession numbers GSE308633 (access token: mzolmweuvpazdgj), GSE308635 (access token: arqluekyxtylbql), GSE308649 (access token: clibaaqyvrevzcn), GSE308692 (access token: kzurmeyidfybtsh), and GSE338983 (access token: anwhacqytronrgb). Additional resources include the SINBA medulloblastoma portal (https://yulab-stjude.shinyapps.io/SINBA_MB), the normal glutamatergic lineage developmental reference portal (https://scminer.stjude.org/study/Glutamatergic-lineage), and the medulloblastoma patient scRNA-seq portal (https://scminer.stjude.org/study/GSE155446). 10x Genomics scRNA-seq data from developing human cerebellum and medulloblastoma patients were obtained from Luo et al.^71^ and are also available via the China National GeneBank Data Base (CNP0002781). Additional medulloblastoma patient 10x scRNA-seq data were downloaded from GEO (GSE155446), and Smart-seq2 scRNA-seq data from GEO (GSE119926).

## CODE AVAILABILITY

SINBA package is available on GitHub (https://github.com/jyyulab/SINBA).

## ACKNOWLEDGEMENTS

We extend our sincere gratitude to Sarah August for her exceptional contributions to this project, including her dedicated efforts in manuscript preparation and revision, her creative input, and her expertise in data presentation. We are also deeply grateful to the following St. Jude Children’s Research Hospital core facilities for their consistent support and expertise: the Center for *In Vivo* Imaging and Therapy (Melissa Johnson and team), the Animal Research Center (Chandra Savage and team), the Flow Cytometry and Cell Sorting Core, and the Comparative Pathology Core. We thank Dr. Anang A. Shelat for his expert assistance in analyzing the high-throughput screening data and Dr. Suzanne J. Baker for her insightful suggestions on the design of our *in vivo* experiments. We are also grateful to Dr. Till Milde for sharing the HDMB03 line and Dr. Q. Richard Lu for generously sharing multiple MB models including CTD. We thank Dr. Martine F. Roussel for her critical review of the manuscript, valuable comments, generous sharing of medulloblastoma models, and assistance in establishing drug-screening protocols. This work is supported, in part, by National Institutes of Health grants R01GM134382 (to J.Y.), U01CA264610 (to J.Y.), and U01CA281868 (to J.Y.), by the COOKIES FOR KIDS CANCER translational award (to J.Y.), by the Big Hope Little Warrior Cancer Foundation, and by the American Lebanese Syrian Associated Charities.

## AUTHOR CONTRIBUTIONS

J.Y. and J.L. conceptualized the study; J.L., X.Y. and M.Z. performed all the experiments with B.B., B.J., W.L., X.F., S.B., J.Y. and S.L.; J.L. performed all the data analysis with X.Y., M.Z., X.D., H.Z., B.J., L.Y., B.B.F.III, A.W. and R.J.; T.C., G.W.R., M.F.R., T.E.M. and A.G. provided relevant intellectual input and edited the manuscript. J.L., X.Y., and M.Z. interpreted the results. J.L., X.Y., M.Z. and J.Y. wrote the manuscript. All the authors reviewed and commented on the manuscript.

## REFERENCES

1. Siegel DA, et al. Pediatric cancer mortality and survival in the United States, 2001-2016. Cancer 126, 4379–4389 (2020).

2. Kool M, et al. Molecular subgroups of medulloblastoma: an international meta-analysis of transcriptome, genetic aberrations, and clinical data of WNT, SHH, Group 3, and Group 4 medulloblastomas. Acta Neuropathol 123, 473–484 (2012).

3. Taylor MD, et al. Molecular subgroups of medulloblastoma: the current consensus. Acta Neuropathol 123, 465–472 (2012).

4. Gajjar A, et al. Risk-adapted craniospinal radiotherapy followed by high-dose chemotherapy and stem-cell rescue in children with newly diagnosed medulloblastoma (St Jude Medulloblastoma-96): long-term results from a prospective, multicentre trial. The Lancet Oncology 7, 813–820 (2006).

5. Gajjar A, et al. Outcomes by Clinical and Molecular Features in Children With Medulloblastoma Treated With Risk-Adapted Therapy: Results of an International Phase III Trial (SJMB03). J Clin Oncol, JCO2001372 (2021).

6. Koschmann C, Bloom K, Upadhyaya S, Geyer JR, Leary SE. Survival After Relapse of Medulloblastoma. J Pediatr Hematol Oncol 38, 269–273 (2016).

7. Sabel M, et al. Relapse patterns and outcome after relapse in standard risk medulloblastoma: a report from the HIT-SIOP-PNET4 study. J Neurooncol 129, 515–524 (2016).

8. Aldape K, et al. Challenges to curing primary brain tumours. Nat Rev Clin Oncol 16, 509–520 (2019).

9. Northcott PA, et al. The whole-genome landscape of medulloblastoma subtypes. Nature 547, 311–317 (2017).

10. Ibrahim YH, et al. PI3K inhibition impairs BRCA1/2 expression and sensitizes BRCA-proficient triple-negative breast cancer to PARP inhibition. Cancer Discov 2, 1036–1047 (2012).

11. Robert C, et al. Improved overall survival in melanoma with combined dabrafenib and trametinib. N Engl J Med 372, 30–39 (2015).

12. DiNardo CD, et al. Azacitidine and Venetoclax in Previously Untreated Acute Myeloid Leukemia. N Engl J Med 383, 617–629 (2020).

13. Turner NC, et al. Palbociclib in Hormone-Receptor-Positive Advanced Breast Cancer. N Engl J Med 373, 209–219 (2015).

14. Mok TSK, et al. Pembrolizumab versus chemotherapy for previously untreated, PD-L1-expressing, locally advanced or metastatic non-small-cell lung cancer (KEYNOTE-042): a randomised, open-label, controlled, phase 3 trial. Lancet 393, 1819–1830 (2019).

15. Larkin J, Hodi FS, Wolchok JD. Combined Nivolumab and Ipilimumab or Monotherapy in Untreated Melanoma. N Engl J Med 373, 1270–1271 (2015).

16. Chen EX, et al. Effect of Combined Immune Checkpoint Inhibition vs Best Supportive Care Alone in Patients With Advanced Colorectal Cancer: The Canadian Cancer Trials Group CO.26 Study. JAMA Oncol 6, 831–838 (2020).

17. Eng C, et al. Atezolizumab with or without cobimetinib versus regorafenib in previously treated metastatic colorectal cancer (IMblaze370): a multicentre, open-label, phase 3, randomised, controlled trial. Lancet Oncol 20, 849–861 (2019).

18. Pei Y, et al. HDAC and PI3K Antagonists Cooperate to Inhibit Growth of MYC-Driven Medulloblastoma. Cancer Cell 29, 311–323 (2016).

19. Endersby R, et al. Small-molecule screen reveals synergy of cell cycle checkpoint kinase inhibitors with DNA-damaging chemotherapies in medulloblastoma. Science Translational Medicine 13, eaba7401 (2021).

20. Morfouace M, et al. Pemetrexed and gemcitabine as combination therapy for the treatment of Group3 medulloblastoma. Cancer Cell 25, 516–529 (2014).

21. Jonchere B, et al. Combination of Ribociclib with BET-Bromodomain and PI3K/mTOR Inhibitors for Medulloblastoma Treatment In Vitro and In Vivo. Mol Cancer Ther 22, 37–51 (2023).

22. Preuer K, Lewis RPI, Hochreiter S, Bender A, Bulusu KC, Klambauer G. DeepSynergy: predicting anti-cancer drug synergy with Deep Learning. Bioinformatics 34, 1538–1546 (2018).

23. Liu W, Wang Z, Liu X, Zeng N, Liu Y, Alsaadi FE. A survey of deep neural network architectures and their applications. Neurocomputing 234, 11–26 (2017).

24. Adam G, Rampasek L, Safikhani Z, Smirnov P, Haibe-Kains B, Goldenberg A. Machine learning approaches to drug response prediction: challenges and recent progress. NPJ Precis Oncol 4, 19 (2020).

25. Jassim A, et al. Gene context drift identifies drug targets to mitigate cancer treatment resistance. Cancer Cell 43, 1608–1621 e1609 (2025).

26. Bansal M, et al. A community computational challenge to predict the activity of pairs of compounds. Nat Biotechnol 32, 1213–1222 (2014).

27. Kuenzi BM, et al. Predicting Drug Response and Synergy Using a Deep Learning Model of Human Cancer Cells. Cancer Cell 38, 672–684 e676 (2020).

28. Robinson G, et al. Novel mutations target distinct subgroups of medulloblastoma. Nature 488, 43–48 (2012).

29. Califano A, Alvarez MJ. The recurrent architecture of tumour initiation, progression and drug sensitivity. Nat Rev Cancer 17, 116–130 (2017).

30. Piovan E, et al. Direct reversal of glucocorticoid resistance by AKT inhibition in acute lymphoblastic leukemia. Cancer Cell 24, 766–776 (2013).

31. Autry RJ, et al. Integrative genomic analyses reveal mechanisms of glucocorticoid resistance in acute lymphoblastic leukemia. Nature Cancer 1, 329–344 (2020).

32. Zeleke TZ, et al. Network-based assessment of HDAC6 activity predicts preclinical and clinical responses to the HDAC6 inhibitor ricolinostat in breast cancer. Nat Cancer 4, 257–275 (2023).

33. Gocho Y, et al. Network-based systems pharmacology reveals heterogeneity in LCK and BCL2 signaling and therapeutic sensitivity of T-cell acute lymphoblastic leukemia. Nature Cancer, (2021).

34. Ianevski A, Giri AK, Aittokallio T. SynergyFinder 2.0: visual analytics of multi-drug combination synergies. Nucleic Acids Res 48, W488–W493 (2020).

35. Yadav B, Wennerberg K, Aittokallio T, Tang J. Searching for Drug Synergy in Complex Dose-Response Landscapes Using an Interaction Potency Model. Comput Struct Biotechnol J 13, 504–513 (2015).

36. Meng F, Xi Y, Huang J, Ayers PW. A curated diverse molecular database of blood-brain barrier permeability with chemical descriptors. Sci Data 8, 289 (2021).

37. Wishart DS, et al. DrugBank 5.0: a major update to the DrugBank database for 2018. Nucleic Acids Res 46, D1074–D1082 (2018).

38. Cheng F, Kovacs IA, Barabasi AL. Network-based prediction of drug combinations. Nat Commun 10, 1197 (2019).

39. Subramanian A, et al. A Next Generation Connectivity Map: L1000 Platform and the First 1,000,000 Profiles. Cell 171, 1437–1452 e1417 (2017).

40. Corsello SM, et al. The Drug Repurposing Hub: a next-generation drug library and information resource. Nat Med 23, 405–408 (2017).

41. Cotto KC, et al. DGIdb 3.0: a redesign and expansion of the drug-gene interaction database. Nucleic Acids Res 46, D1068–D1073 (2018).

42. Cannon M, et al. DGIdb 5.0: rebuilding the drug-gene interaction database for precision medicine and drug discovery platforms. Nucleic Acids Res 52, D1227–D1235 (2024).

43. Griffith M, et al. DGIdb: mining the druggable genome. Nat Methods 10, 1209–1210 (2013).

44. Khatamian A, Paull EO, Califano A, Yu J. SJARACNe: a scalable software tool for gene network reverse engineering from big data. Bioinformatics 35, 2165–2166 (2019).

45. Northcott PA, et al. Subgroup-specific structural variation across 1,000 medulloblastoma genomes. Nature 488, 49–56 (2012).

46. de Bont JM, et al. Differential expression and prognostic significance of SOX genes in pediatric medulloblastoma and ependymoma identified by microarray analysis. Neuro Oncol 10, 648–660 (2008).

47. Ashburner M, et al. Gene Ontology: tool for the unification of biology. Nature Genetics 25, 25–29 (2000).

48. Dong X, et al. NetBID2 provides comprehensive hidden driver analysis. Nat Commun 14, 2581 (2023).

49. Cavalli FMG, et al. Intertumoral Heterogeneity within Medulloblastoma Subgroups. Cancer Cell 31, 737–754 e736 (2017).

50. Archer TC, et al. Proteomics, Post-translational Modifications, and Integrative Analyses Reveal Molecular Heterogeneity within Medulloblastoma Subgroups. Cancer Cell 34, 396–410 e398 (2018).

51. Forget A, et al. Aberrant ERBB4-SRC Signaling as a Hallmark of Group 4 Medulloblastoma Revealed by Integrative Phosphoproteomic Profiling. Cancer Cell 34, 379–395 e377 (2018).

52. Scrucca L. GA: A Package for Genetic Algorithms in R. Journal of Statistical Software 53, 1–37 (2013).

53. Kort A, Durmus S, Sparidans RW, Wagenaar E, Beijnen JH, Schinkel AH. Brain and Testis Accumulation of Regorafenib is Restricted by Breast Cancer Resistance Protein (BCRP/ABCG2) and P-glycoprotein (P-GP/ABCB1). Pharm Res 32, 2205–2216 (2015).

54. Jiang J, et al. Regorafenib induces lethal autophagy arrest by stabilizing PSAT1 in glioblastoma. Autophagy 16, 106–122 (2020).

55. Zeiner PS, et al. Regorafenib CSF Penetration, Efficacy, and MRI Patterns in Recurrent Malignant Glioma Patients. J Clin Med 8, (2019).

56. Freeman BB, 3rd, Yang L, Rankovic Z. Practical approaches to evaluating and optimizing brain exposure in early drug discovery. Eur J Med Chem 182, 111643 (2019).

57. de Gooijer MC, Zhang P, Weijer R, Buil LCM, Beijnen JH, van Tellingen O. The impact of P-glycoprotein and breast cancer resistance protein on the brain pharmacokinetics and pharmacodynamics of a panel of MEK inhibitors. Int J Cancer 142, 381–391 (2018).

58. He C, et al. Patient-derived models recapitulate heterogeneity of molecular signatures and drug response in pediatric high-grade glioma. Nat Commun 12, 4089 (2021).

59. Klaeger S, et al. The target landscape of clinical kinase drugs. Science 358, (2017).

60. Kanehisa M, Goto S. KEGG: kyoto encyclopedia of genes and genomes. Nucleic Acids Res 28, 27–30 (2000).

61. Ghandi M, et al. Next-generation characterization of the Cancer Cell Line Encyclopedia. Nature 569, 503–508 (2019).

62. Pilarczyk M, et al. Connecting omics signatures and revealing biological mechanisms with iLINCS. Nat Commun 13, 4678 (2022).

63. Smith KS, et al. Patient-derived orthotopic xenografts of pediatric brain tumors: a St. Jude resource. Acta Neuropathol 140, 209–225 (2020).

64. McLeod C, et al. St. Jude Cloud: A Pediatric Cancer Genomic Data-Sharing Ecosystem. Cancer Discov 11, 1082–1099 (2021).

65. Padovani L, Horan G, Ajithkumar T. Radiotherapy Advances in Paediatric Medulloblastoma Treatment. Clin Oncol (R Coll Radiol) 31, 171–181 (2019).

66. Hill RM, et al. Time, pattern, and outcome of medulloblastoma relapse and their association with tumour biology at diagnosis and therapy: a multicentre cohort study. The Lancet Child & Adolescent Health 4, 865–874 (2020).

67. Hovestadt V, et al. Resolving medulloblastoma cellular architecture by single-cell genomics. Nature, (2019).

68. Riemondy KA, et al. Neoplastic and immune single-cell transcriptomics define subgroup-specific intra-tumoral heterogeneity of childhood medulloblastoma. Neuro Oncol 24, 273–286 (2022).

69. Hendrikse LD, et al. Failure of human rhombic lip differentiation underlies medulloblastoma formation. Nature 609, 1021–1028 (2022).

70. Smith KS, et al. Unified rhombic lip origins of group 3 and group 4 medulloblastoma. Nature 609, 1012–1020 (2022).

71. Luo Z, et al. Human fetal cerebellar cell atlas informs medulloblastoma origin and oncogenesis. bioRxiv, 2022.2008.2017.504304 (2022).

72. Gibson P, et al. Subtypes of medulloblastoma have distinct developmental origins. Nature 468, 1095–1099 (2010).

73. Williams JS, et al. Patient-derived pediatric brain tumor orthotopic xenografts and tumor organoids faithfully recapitulate primary tumors. Sci Adv 12, eaea4966 (2026).

74. Erickson AW, et al. Mapping the developmental profile of ventricular zone-derived neurons in the human cerebellum. Proc Natl Acad Sci U S A 122, e2415425122 (2025).

75. Sepp M, et al. Cellular development and evolution of the mammalian cerebellum. Nature, (2024).

76. Aldinger KA, et al. Spatial and cell type transcriptional landscape of human cerebellar development. Nat Neurosci 24, 1163–1175 (2021).

77. Lu S, et al. Transcriptome size matters for single-cell RNA-seq normalization and bulk deconvolution. Nat Commun 16, 1246 (2025).

78. Luo Z, et al. Genomic and Transcriptomic Analyses Reveals ZNF124 as a Critical Regulator in Highly Aggressive Medulloblastomas. Front Cell Dev Biol 9, 634056 (2021).

79. Pan Q, et al. scMINER: a mutual information-based framework for clustering and hidden driver inference from single-cell transcriptomics data. Nat Commun 16, 4305 (2025).

80. Luo Z, et al. Loss of phosphatase CTDNEP1 potentiates aggressive medulloblastoma by triggering MYC amplification and genomic instability. Nat Commun 14, 762 (2023).

81. Bae J, et al. Targeting LAG3/GAL-3 to overcome immunosuppression and enhance anti-tumor immune responses in multiple myeloma. Leukemia 36, 138–154 (2022).

82. Blum JS, Wearsch PA, Cresswell P. Pathways of antigen processing. Annu Rev Immunol 31, 443–473 (2013).

83. Chen YH, et al. Regorafenib enhances M1/M2 macrophage polarization by inhibiting the secretion of plasminogen activator inhibitor-1 in head and neck squamous cell carcinoma. Life Sci 358, 123147 (2024).

84. Allen F, et al. CCL3 augments tumor rejection and enhances CD8(+) T cell infiltration through NK and CD103(+) dendritic cell recruitment via IFNgamma. Oncoimmunology 7, e1393598 (2018).

85. Kang TG, Park HJ, Moon J, Lee JH, Ha SJ. Enriching CCL3 in the Tumor Microenvironment Facilitates T cell Responses and Improves the Efficacy of Anti-PD-1 Therapy. Immune Netw 21, e23 (2021).

86. Eberlein J, et al. Chemokine Signatures of Pathogen-Specific T Cells I: Effector T Cells. J Immunol 205, 2169–2187 (2020).

87. Wang BL, et al. CXCR6 is required for antitumor efficacy of intratumoral CD8 T cell. Journal for Immunotherapy of Cancer 9, (2021).

88. Di Pilato M, et al. CXCR6 positions cytotoxic T cells to receive critical survival signals in the tumor microenvironment. Cell 184, 4512–+ (2021).

89. Muthuswamy R, et al. CXCR6 by increasing retention of memory CD8 T cells in the ovarian tumor microenvironment promotes immunosurveillance and control of ovarian cancer. Journal for Immunotherapy of Cancer 9, (2021).

90. Seo YD, et al. Mobilization of CD8+ T Cells via CXCR4 Blockade Facilitates PD-1 Checkpoint Therapy in Human Pancreatic Cancer. Clinical Cancer Research 25, 3934–3945 (2019).

91. Liu SY, Zhang W, Liu K, Wang YC. CD160 expression on CD8 T cells is associated with active effector responses but limited activation potential in pancreatic cancer. Cancer Immunol Immun 69, 789–797 (2020).

92. Duhen T, et al. Co-expression of CD39 and CD103 identifies tumor-reactive CD8 T cells in human solid tumors. Nature Communications 9, (2018).

93. Lamichhane R, et al. TCR- or Cytokine-Activated CD8(+) Mucosal-Associated Invariant T Cells Are Rapid Polyfunctional Effectors That Can Coordinate Immune Responses. Cell Rep 28, 3061–3076 e3065 (2019).

94. Jin SQ, et al. Inference and analysis of cell-cell communication using CellChat. Nature Communications 12, (2021).

95. Mestas J, Hughes CC. Of mice and not men: differences between mouse and human immunology. J Immunol 172, 2731–2738 (2004).

96. Olson B, Li Y, Lin Y, Liu ET, Patnaik A. Mouse Models for Cancer Immunotherapy Research. Cancer Discov 8, 1358–1365 (2018).

97. Ou DL, et al. Regorafenib enhances antitumor immunity via inhibition of p38 kinase/Creb1/Klf4 axis in tumor-associated macrophages. J Immunother Cancer 9, (2021).

98. Kanikarla Marie P, et al. Autologous humanized mouse models to study combination and single-agent immunotherapy for colorectal cancer patient-derived xenografts. Front Oncol 12, 994333 (2022).

99. Kozareva V, et al. A transcriptomic atlas of mouse cerebellar cortex comprehensively defines cell types. Nature 598, 214–219 (2021).

100. Huang L, et al. Systems biology-based drug repositioning identifies digoxin as a potential therapy for groups 3 and 4 medulloblastoma. Science Translational Medicine 10, (2018).

101. Hovestadt V, Ayrault O, Swartling FJ, Robinson GW, Pfister SM, Northcott PA. Medulloblastomics revisited: biological and clinical insights from thousands of patients. Nat Rev Cancer 20, 42–56 (2020).

102. Visvanathan A, et al. Early rhombic lip Protogenin(+ve) stem cells in a human-specific neurovascular niche initiate and maintain group 3 medulloblastoma. Cell 187, 4733–4750 e4726 (2024).

103. Jessa S, et al. Stalled developmental programs at the root of pediatric brain tumors. Nat Genet 51, 1702–1713 (2019).

104. Xie X, et al. Converging mechanism of UM171 and KBTBD4 neomorphic cancer mutations. Nature 639, 241–249 (2025).

105. Corcoran RB, et al. Combined BRAF and MEK Inhibition With Dabrafenib and Trametinib in BRAF V600-Mutant Colorectal Cancer. J Clin Oncol 33, 4023–4031 (2015).

106. Wu CS, et al. Curcumin functions as a MEK inhibitor to induce a synthetic lethal effect on KRAS mutant colorectal cancer cells receiving targeted drug regorafenib. J Nutr Biochem 74, 108227 (2019).

107. Lau DK, et al. Rapid Resistance of FGFR-driven Gastric Cancers to Regorafenib and Targeted FGFR Inhibitors can be Overcome by Parallel Inhibition of MEK. Mol Cancer Ther 20, 704–715 (2021).

108. Adamopoulos C, et al. Exploiting Allosteric Properties of RAF and MEK Inhibitors to Target Therapy-Resistant Tumors Driven by Oncogenic BRAF Signaling. Cancer Discov, (2021).

109. MacDonald TJ, et al. Expression profiling of medulloblastoma: PDGFRA and the RAS/MAPK pathway as therapeutic targets for metastatic disease. Nature Genetics 35, 287–287 (2003).

110. Abeysundara N, et al. Metastatic medulloblastoma remodels the local leptomeningeal microenvironment to promote further metastatic colonization and growth. Nat Cell Biol, (2025).

111. da Cunha Jaeger M, et al. HDAC and MAPK/ERK Inhibitors Cooperate To Reduce Viability and Stemness in Medulloblastoma. J Mol Neurosci 70, 981–992 (2020).

112. Bakhshinyan D, et al. Temporal profiling of therapy resistance in human medulloblastoma identifies novel targetable drivers of recurrence. Sci Adv 7, eabi5568 (2021).

113. Li S, et al. Liposomal Honokiol induces ROS-mediated apoptosis via regulation of ERK/p38-MAPK signaling and autophagic inhibition in human medulloblastoma. In: Signal Transduct Target Ther). 20220221 edn (2022).

114. Lombardi G, et al. Regorafenib compared with lomustine in patients with relapsed glioblastoma (REGOMA): a multicentre, open-label, randomised, controlled, phase 2 trial. The Lancet Oncology 20, 110–119 (2019).

115. Weiss BD, et al. NF106: A Neurofibromatosis Clinical Trials Consortium Phase II Trial of the MEK Inhibitor Mirdametinib (PD-0325901) in Adolescents and Adults With NF1-Related Plexiform Neurofibromas. J Clin Oncol, JCO2002220 (2021).

116. Banerjee A, et al. A phase I study of AZD6244 in children with recurrent or refractory low-grade gliomas: A Pediatric Brain Tumor Consortium report. Journal of Clinical Oncology 32, (2014).

117. Andrew N, et al. Phase 1 Study of Fluvastatin-Celecoxib Combination in Children with Relapsing/Refractory Optico-Chiasmatic Low-Grade Glioma or High-Grade Gliomas (Fluvabrex): Final Results. Neuro-Oncology 22, 305–306 (2020).

118. Ebert PJR, et al. MAP Kinase Inhibition Promotes T Cell and Anti-tumor Activity in Combination with PD-L1 Checkpoint Blockade. Immunity 44, 609–621 (2016).

119. Liu J, et al. Immunomodulatory effects of regorafenib: Enhancing the efficacy of anti-PD-1/PD-L1 therapy. Front Immunol 13, 992611 (2022).

120. Bentayebi K, Louati S, Suwan K, Eljaoudi R, Hajitou A. New developments and prospects for drug delivery in medulloblastoma. Mol Ther Oncol 34, 201186 (2026).

121. Margolin AA, et al. ARACNE: an algorithm for the reconstruction of gene regulatory networks in a mammalian cellular context. BMC Bioinformatics 7 **Suppl 1**, S7 (2006).

122. Ritz C, Baty F, Streibig JC, Gerhard D. Dose-Response Analysis Using R. PLoS One 10, e0146021 (2015).

123. Zheng S, et al. SynergyFinder Plus: Toward Better Interpretation and Annotation of Drug Combination Screening Datasets. Genomics Proteomics Bioinformatics 20, 587–596 (2022).

124. Milde T, et al. HD-MB03 is a novel Group 3 medulloblastoma model demonstrating sensitivity to histone deacetylase inhibitor treatment. J Neurooncol 110, 335–348 (2012).

125. Bandopadhayay P, et al. BET bromodomain inhibition of MYC-amplified medulloblastoma. Clin Cancer Res 20, 912–925 (2014).

126. Kawauchi D, et al. A mouse model of the most aggressive subgroup of human medulloblastoma. Cancer Cell 21, 168–180 (2012).

127. Pribnow A, et al. Combination of Ribociclib and Gemcitabine for the Treatment of Medulloblastoma. Mol Cancer Ther 21, 1306–1317 (2022).

128. Xie J, et al. ATM inhibition enhances the efficacy of radiation across distinct molecular subgroups of pediatric high-grade glioma. Neuro Oncol 25, 1828–1841 (2023).

129. Veo B, et al. Single-cell multi-omics identifies metabolism-linked epigenetic reprogramming as a driver of therapy-resistant medulloblastoma. Nat Commun 16, 10470 (2025).

130. Street K, et al. Slingshot: cell lineage and pseudotime inference for single-cell transcriptomics. BMC Genomics 19, 477 (2018).

131. Qiu X, et al. Reversed graph embedding resolves complex single-cell trajectories. Nat Methods 14, 979–982 (2017).

132. Hao Y, et al. Dictionary learning for integrative, multimodal and scalable single-cell analysis. Nat Biotechnol 42, 293–304 (2024).

