## Supplemental_Figures for "Hidden-driver inference reveals synergistic brain-penetrant therapies for medulloblastoma"

Liu, et al.

**Supplementary Figures 1-11**

**Supplementary Tables 1-16**

**Supplementary Figures**

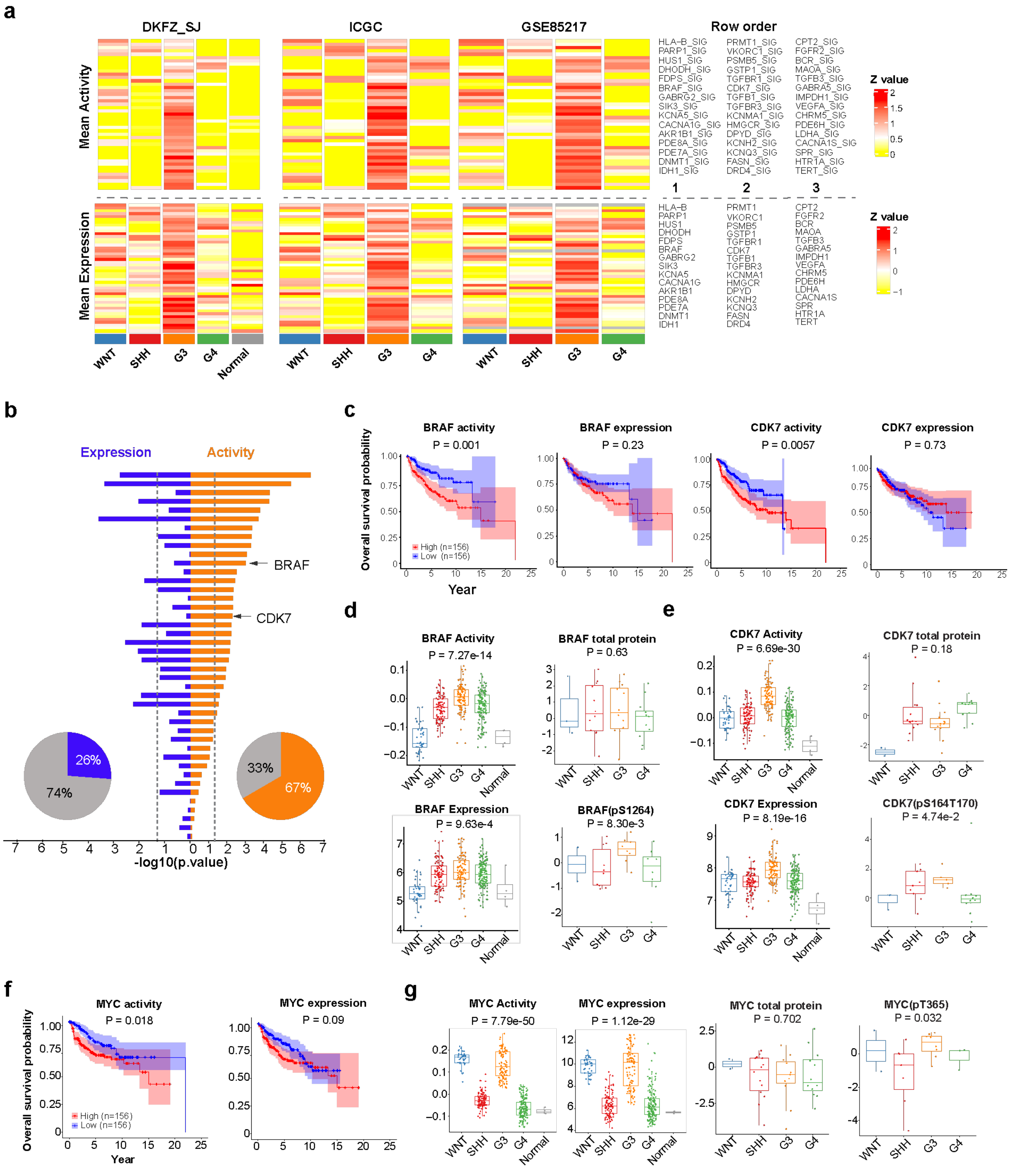


**Figure S1. Druggable hidden drivers in high-risk medulloblastoma.** **a** Heatmap displaying druggable hidden driver activities and expressions in three medulloblastoma cohorts. **b** Druggable hidden driver genes identified by the SINBA algorithm were evaluated for association with overall survival in the GSE85217 cohort. MB Patients were stratified by high (>75th percentile) versus low (<25th percentile) gene expression or activity. Kaplan–Meier survival analysis; p values calculated by log-rank test. **c** Kaplan–Meier curves showing overall survival of MB patients stratified by BRAF or CDK7 activity or expression. **d** BRAF activity, mRNA expression, total protein and phosphorylated BRAF (S1264) levels across MB subgroups and normal samples. **e** CDK7 activity, mRNA expression, total protein and phosphorylated CDK7 (S164/T170) levels across MB subgroups and normal samples. **f** Kaplan–Meier curves showing overall survival of MB patients stratified by MYC activity or expression. **g** MYC activity, mRNA expression, total protein and phosphorylated MYC (T365) levels across MB subgroups and normal samples. For (**d**), (**e**), and (**g**), unpaired two-sided t tests were performed between G3 MB and other subgroups. For (**c**) and (**f**), P value was calculated by log-rank test.

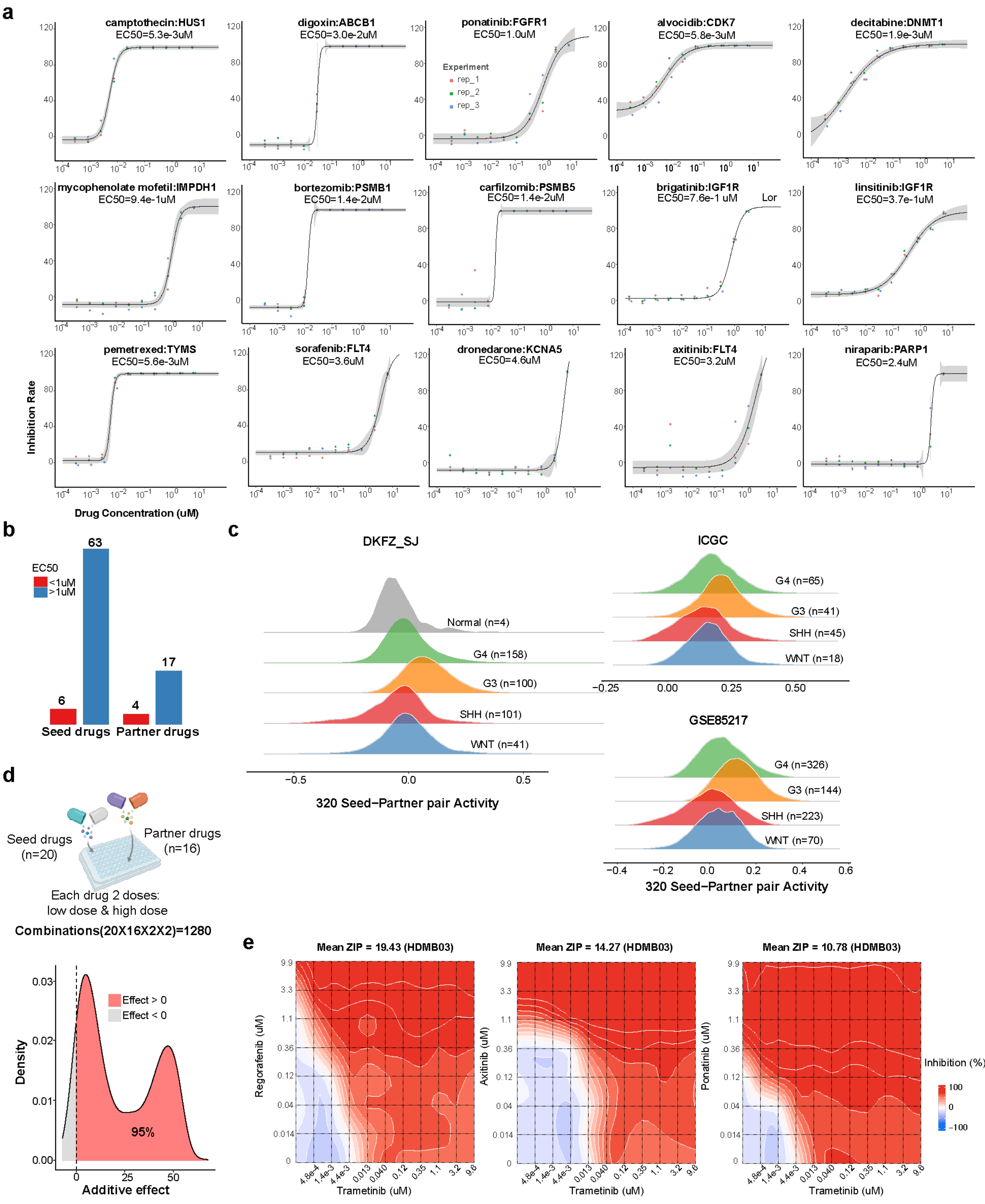


**Figure S2. SINBA-guided multi-step drug screening identifies potential therapeutic combinations for high-risk medulloblastoma. a** Seed and partner drugs with half-maximal effective concentrations ($EC_{50}$) below 10 μM. **b** Summary of HTS results from the seed drug and partner drug libraries. **c** Ridge plots showing the distribution of 320 seed–partner activity scores (20 seeds × 16 partners) in the three MB patient cohorts stratified by molecular subtype. **d** Overview of two-dose combination screening outcomes. **e** Representative contour plots showing HDMB03 cell growth inhibition across varying concentrations of trametinib, axitinib or ponatinib in combination with regorafenib.

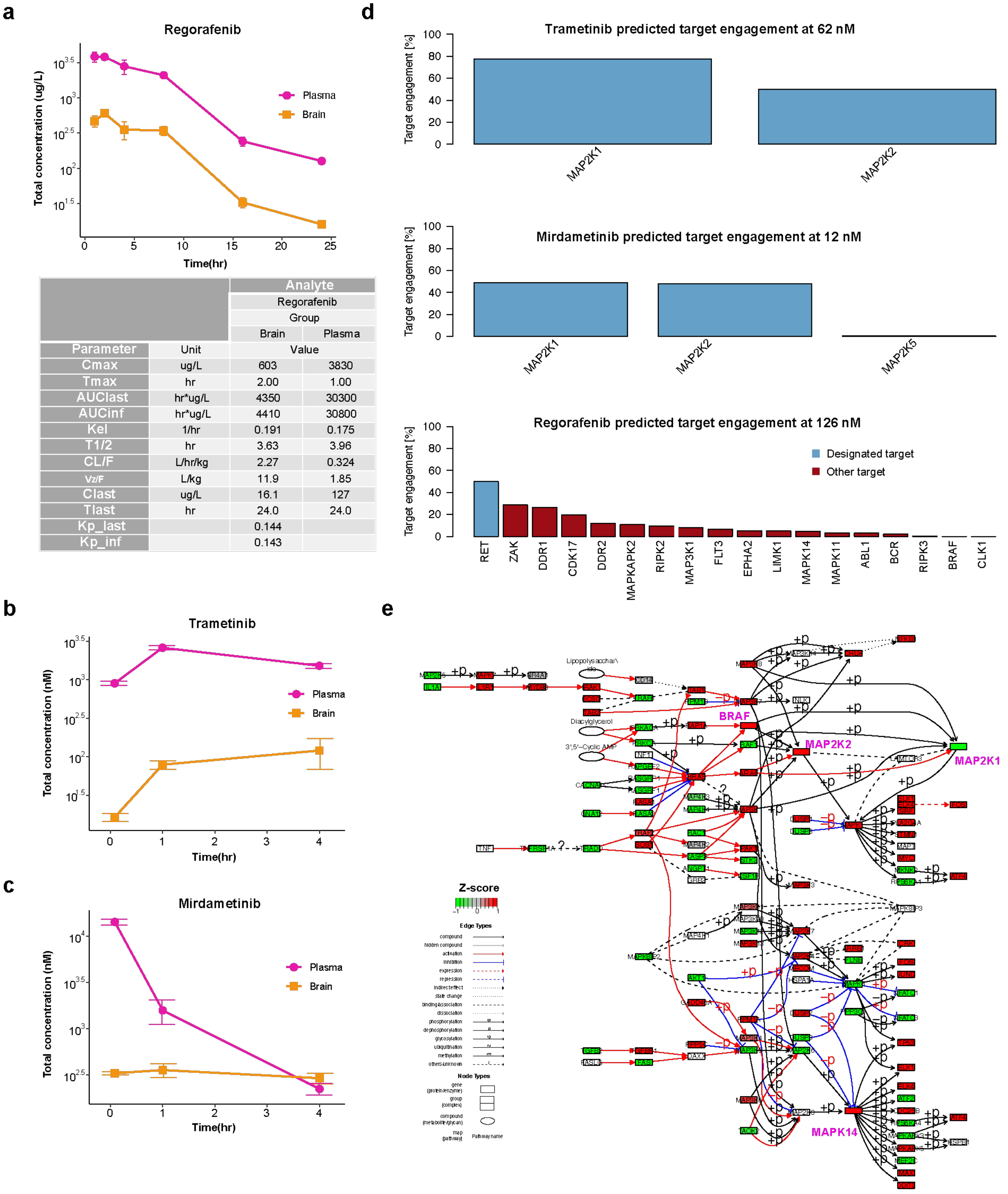


| | | Analyte | |
|---|---|---|---|
| | | Regorafenib | |
| | | Group | |
| | | Brain | Plasma |
| Parameter | Unit | Value | |
| Cmax | ug/L | 603 | 3830 |
| Tmax | hr | 2.00 | 1.00 |
| AUClast | hr*ug/L | 4350 | 30300 |
| AUCinf | hr*ug/L | 4410 | 30800 |
| Kel | 1/hr | 0.191 | 0.175 |
| T1/2 | hr | 3.63 | 3.96 |
| CL/F | L/hr/kg | 2.27 | 0.324 |
| Vz/F | L/kg | 11.9 | 1.85 |
| Clast | ug/L | 16.1 | 127 |
| Tlast | hr | 24.0 | 24.0 |
| Kp_last | | 0.144 | |
| Kp_inf | | 0.143 | |

**Figure S3. Pharmacokinetic profile of the MEK inhibitors and regorafenib. a** Pharmacokinetic profile of regorafenib: plasma and brain concentrations measured over 24 hours post-dose with the statistical analysis of regorafenib plasma and brain concentration–time (Ct) profiles. **b-c** Pharmacokinetic profiles of trametinib and mirdametinib: plasma and brain concentrations measured over 4 hours post-dose. **d** Chemical proteomic profiling of clinical kinase inhibitors, showing predicted interaction targets of trametinib, mirdametinib, and regorafenib. **e** Comparative analysis of driver activity within the KEGG MAPK signaling cascade between G3 and normal samples in the DKFZ_SJ cohort. Nodes are color-coded according to Z-score (normalized p value) differentials reflecting G3 versus normal cerebellum. Genes targeted by trametinib and regorafenib are accentuated in magenta. For (**a**), (**b**), and (**c**), each time point data are represented as mean ± SEM.

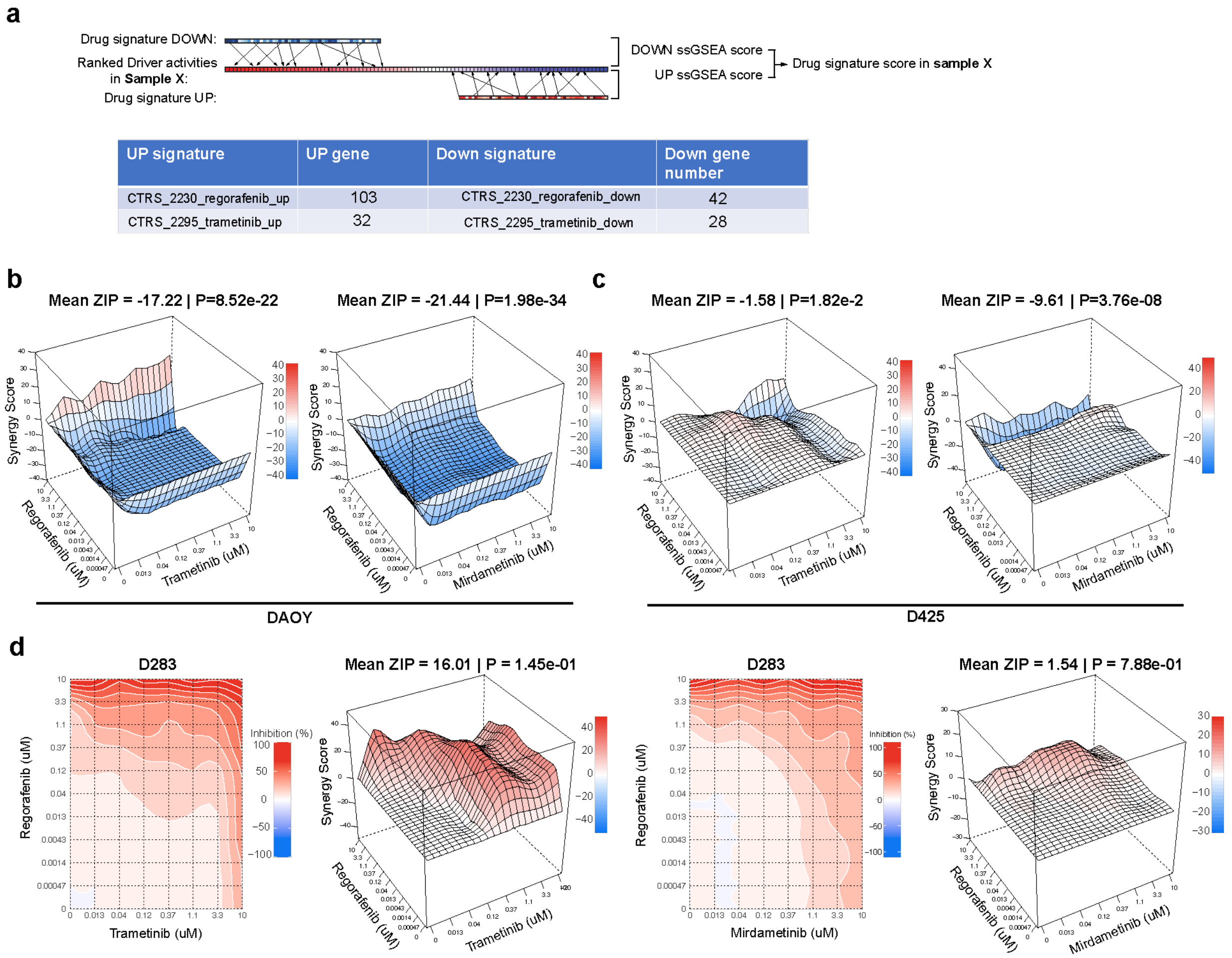


**Figure S4. Drug sensitivity and synergy analysis for MEK inhibitor and regorafenib combinations in medulloblastoma models.** **a** Schematic representation of the drug sensitivity calculation framework, leveraging CNS-specific drug signatures from the iLINCS database to identify responsive medulloblastoma models. Refer to the Methods section for further details. **b** Surface plots of ZIP synergy score landscapes for trametinib plus regorafenib (left) and mirdametinib plus regorafenib (right) in the DAOY cell line. **c** Surface plots of ZIP synergy score landscapes for trametinib plus regorafenib (left) and mirdametinib plus regorafenib (right) in the D425 cell line. **d** Contour plots of cell viability and surface plots of ZIP synergy score landscapes for trametinib plus regorafenib (left) and mirdametinib plus regorafenib (right) in D283 cells.

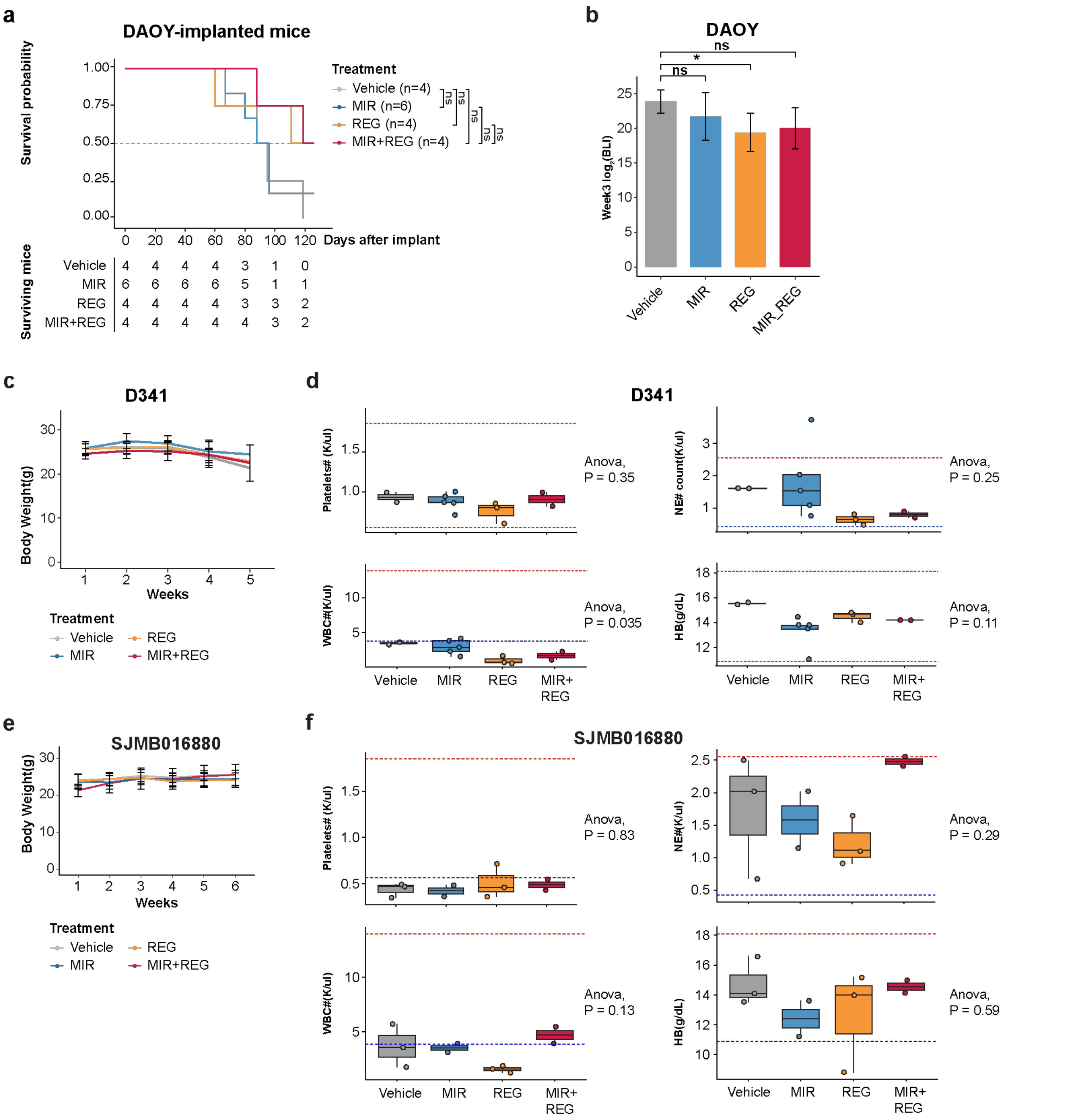


**Figure S5. The MEKi + regorafenib combination lacks synergistic efficacy in the SHH model DAOY and shows no significant toxicity in mice. a** Kaplan–Meier survival curves for mice implanted with DAOY cells and treated as in **Fig. 3a**. Group sizes (n) are indicated in the risk table; log-rank test: ns, not significant. **b** Quantified bioluminescence imaging (BLI) for each treatment group in the DAOY model at week 3 post-treatment. Data are mean ± SEM. **c** Weekly body weight changes in D341-bearing mice after surgery. Data are represented as mean ± SEM. **d** Endpoint toxicity assessment by complete blood counts (n = 2–4 per group) in the D341 study; dashed lines indicate normal reference ranges. **e** Weekly body weight changes in SJMB016880-bearning mice after surgery. Data are represented as mean ± SEM. **f** Endpoint toxicity assessment by complete blood counts (n = 2–4 per group) in the SJMB016880 study. For (**b**) unpaired two-sided t tests were used. *P < 0.05, **P < 0.01, ***P < 0.001; ns, not significant.For (**d**) and (**f**), one-way ANOVA was performed among treatment groups.

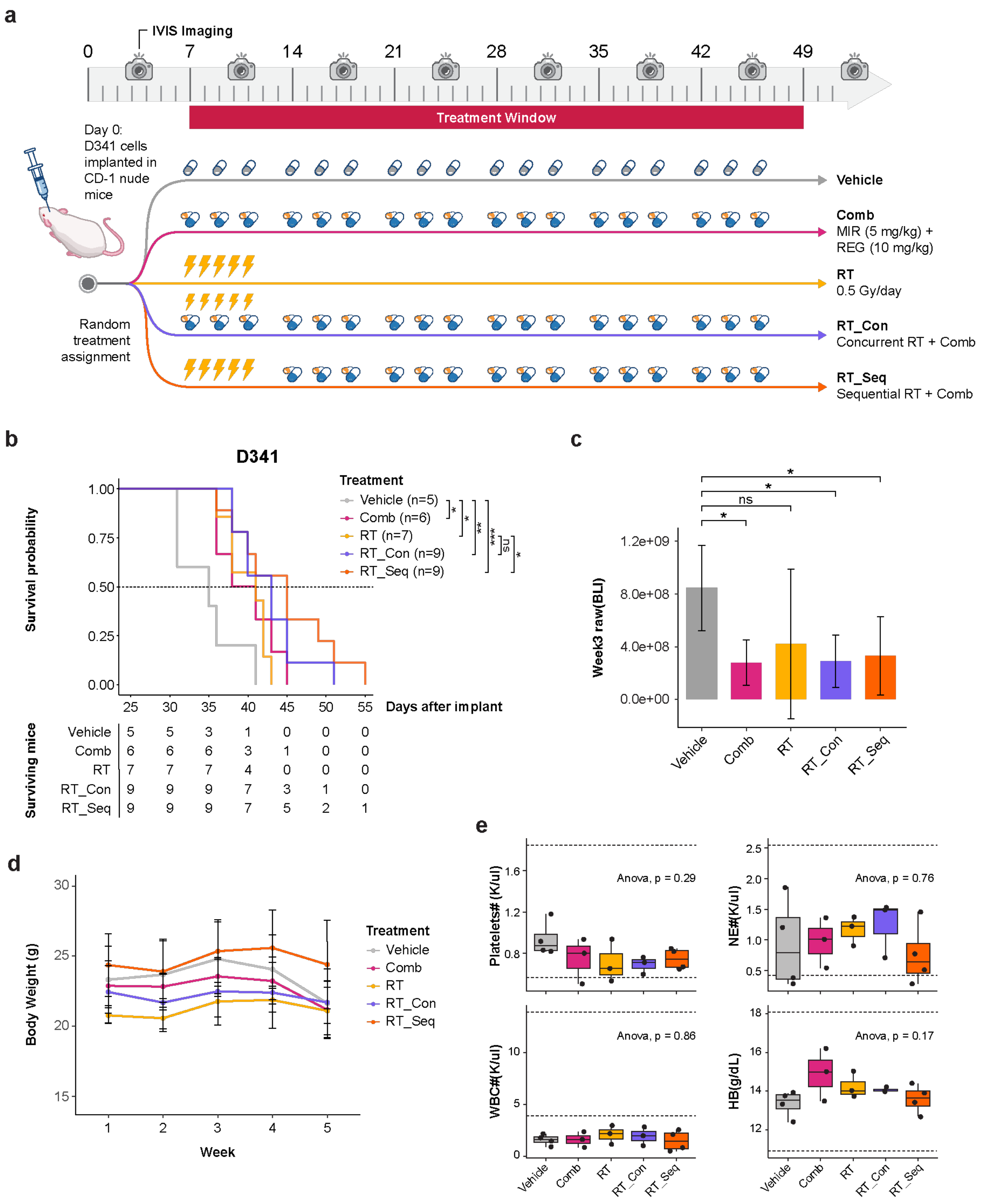


**Figure S6. Evaluating the combination of mirdametinib, regorafenib, and radiation therapy in the D341 Model. a** Schematic of experimental design for drug administration and bioluminescence imaging (IVIS) in CD-1 nude mice. Mice with intracranial D341 cell implants were randomized into four treatment groups: vehicle (p.o.), drug combination (Comb: mirdametinib at 5 mg/kg p.o. and Regorafenib at 10 mg/kg p.o.), radiation therapy (RT: 0.5 Gy/day for 5 consecutive days), concurrent RT + drug combination (RT_Con), or sequential RT + drug combination (RT_Seq). Treatments commenced one week post-implantation and continued for six weeks. **b** Kaplan–Meier survival curves for mice treated as described in (**a**). Group sizes (n) are indicated in the risk table; log-rank test results: *$P < 0.05$, **$P < 0.01$, ***$P < 0.001$. **c** Quantified bioluminescence imaging (BLI) measurements for each treatment group in the D341 model at week 3 post-treatment enrollment. Data are represented as mean ± SEM. Unpaired two-sided t tests were used. *$P < 0.05$, ns, not significant. **d** Weekly body weight changes in D341-bearing mice after surgery. Data are represented as mean ± SEM. **e** Endpoint toxicity assessment by complete blood counts (n = 2–4 per group) in the D341 study as shown in (**a**).

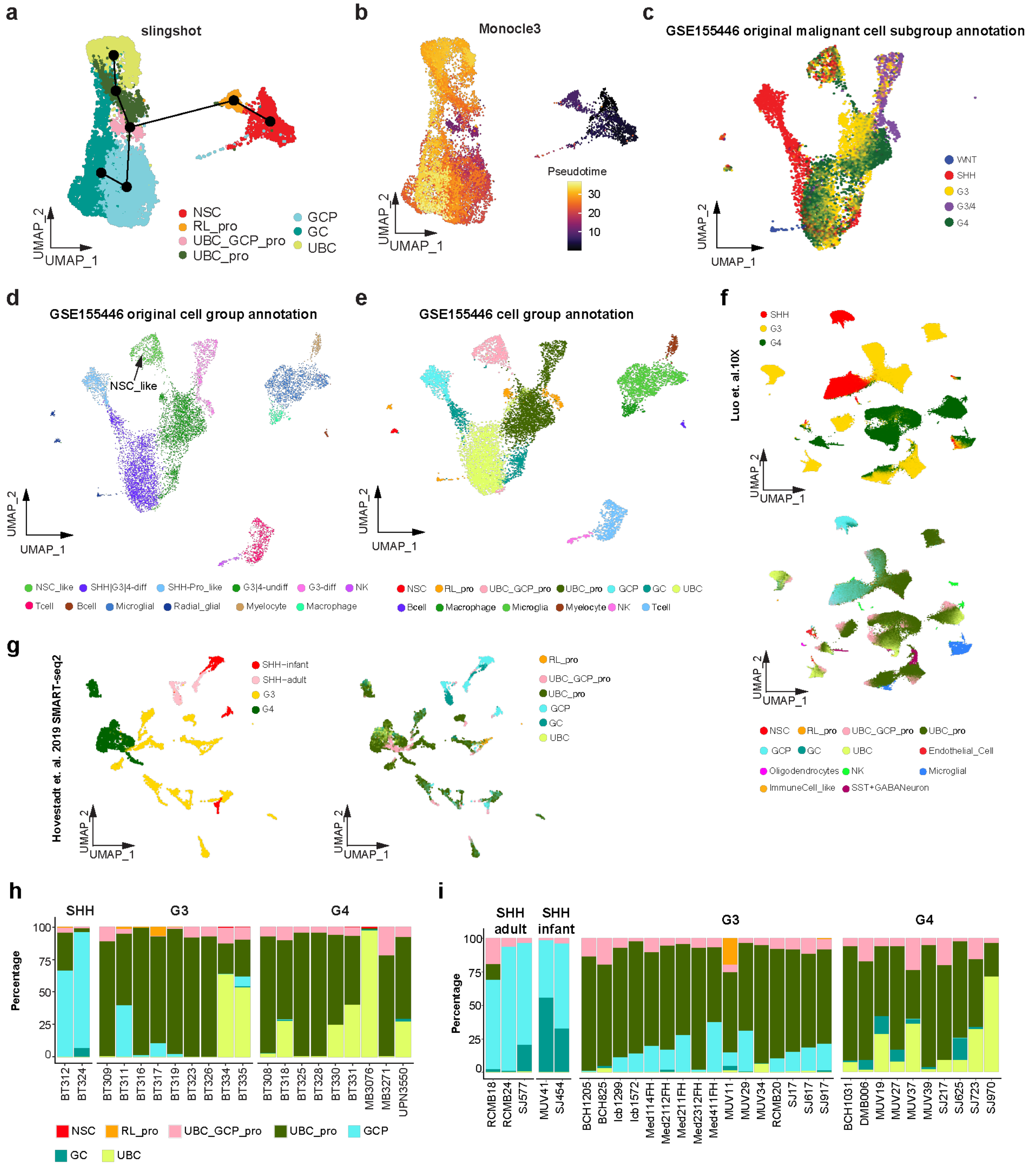


86

87 **Figure S7. Enrichment of UBC_GCP_pro and UBC_pro like tumor cells in G3 and G4 MB revealed**

88 **by scRNA-seq. a** UMAP plot showing developmental trajectories of normal glutamatergic lineage neurons

inferred by Slingshot. **b** UMAP plot of pseudotime trajectories generated by Monocle3, with cells colored by inferred developmental time. **c-d** UMAP plots displaying medulloblastoma tumor subgroup and cell type annotations from the original study (GSE156053; Riemondy et al.). **e** UMAP plot of tumor cell state and normal cell types (GSE156053). **f-g** UMAP plots of two medulloblastoma patient scRNA-seq datasets, annotated by MB subgroup or cell type. **h-i** Proportional distribution of tumor cell states in individual medulloblastoma patients.

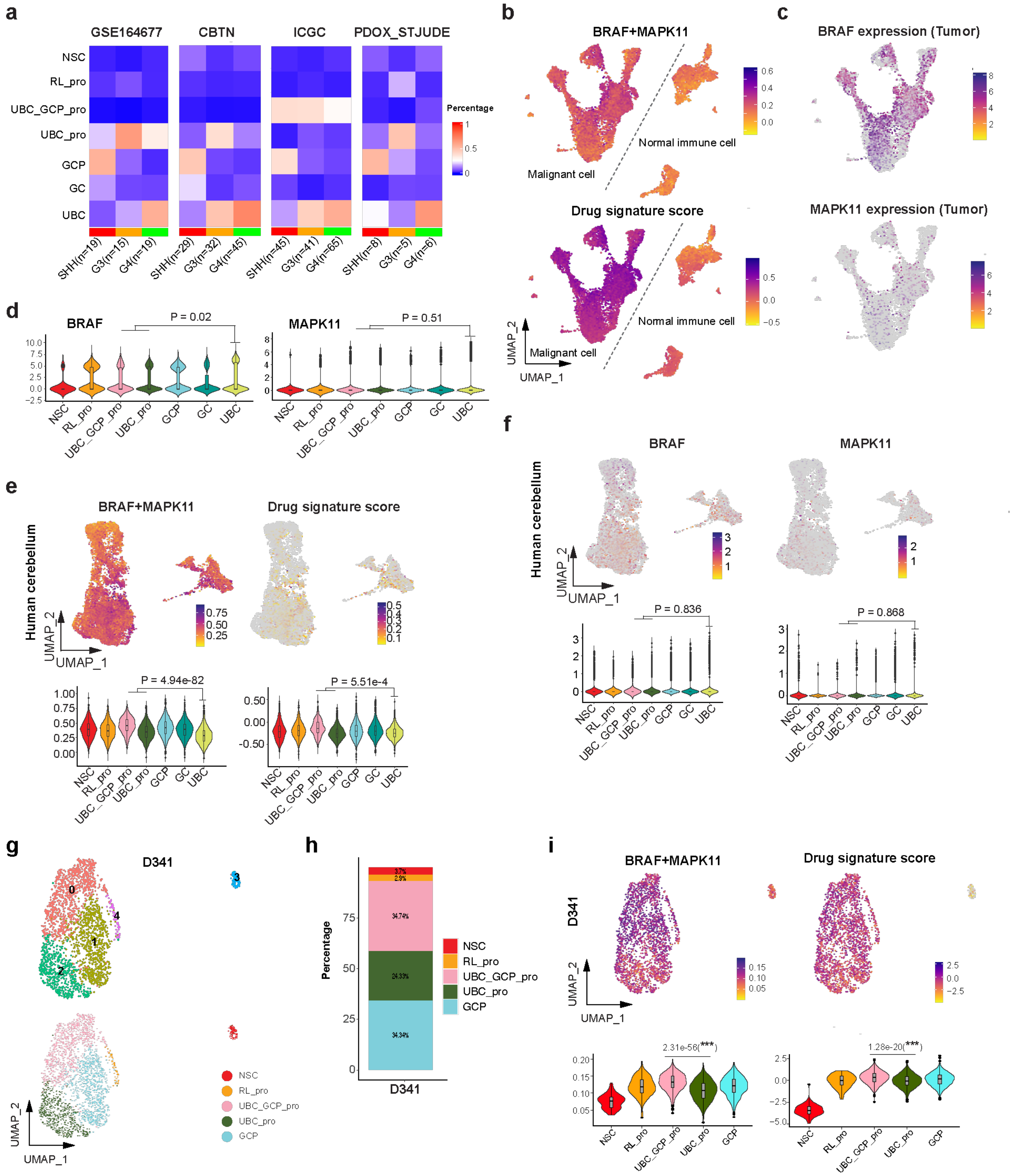


**Figure S8. UBC_GCP_Pro like tumor cells exhibiting the highest driver activities and drug signature scores.** a Transcriptomic deconvolution of RNA-seq profiles via the ReDeconv algorithm across three independent medulloblastoma patient cohorts and a PDOX dataset, stratified by cellular subpopulations as delineated in Figure 5A. **b** Feature plot of driver combination activities and drug signature scores in malignant cells and normal immune cells (GSE156053). **c-d** Feature and violin plots of BRAF and MAPK11 expression in tumor cells (GSE156053). **e** Feature and violin plots of driver combination activity and drug signature scores in the normal reference. **f** Feature and violin plots of BRAF and MAPK11 expression in the normal reference. **g** UMAP of D341 cell clusters and tumor cell states. **h** Proportional distribution of tumor cell states in D341. **i** Feature and violin plots showing driver combination activities and predicted drug sensitivity scores across tumor cell sate. For (**d**), (**e**), and (**f**), unpaired two-sided t tests were used to compare progenitor-like UBC_GCP_pro and GCP_pro cells with mature-like UBC cells. For (**i**), unpaired two-sided t tests were used to compare UBC_GCP_pro with UBC_pro cells.

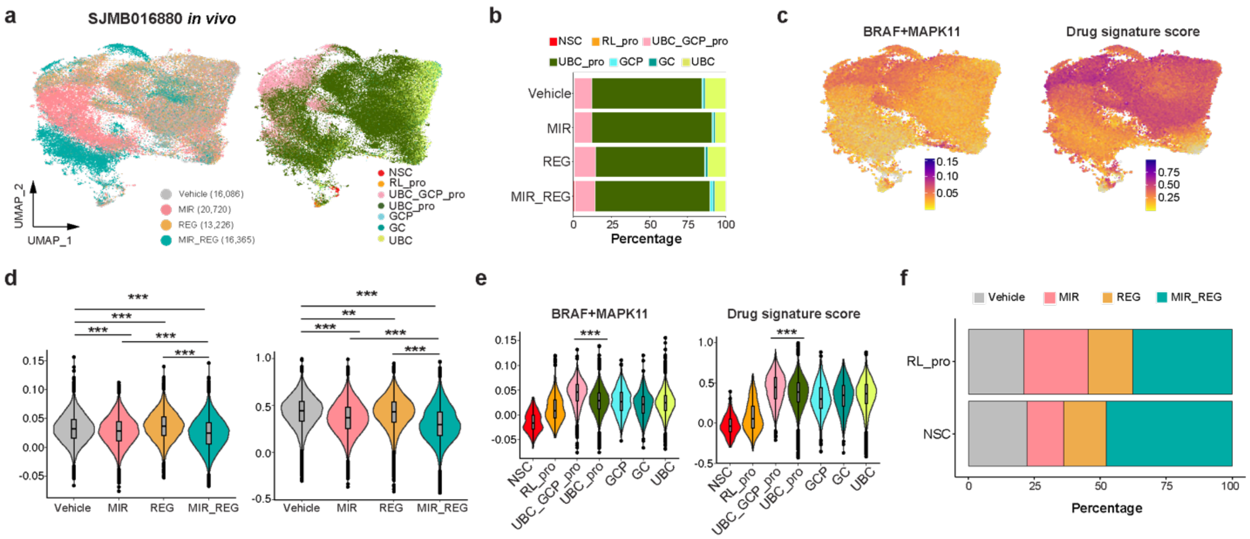


**Figure S9. MEK inhibitor and regorafenib combination therapy targets UBC_GCP common progenitor and UBC progenitor-like tumor cells in the PDX model.** **a** UMAP visualization of orthotopically implanted SJMB016880 tumor cells collected from treated CD-1 mice and annotated with tumor cell states using a normal developmental reference. **b** Proportional distribution of tumor cell states across treatment groups. **c-e** Single-cell feature plots **(c)** and violin plots **(d, e)** showing driver activities and drug sensitivity scores across treatment groups **(d)** and predicted cell states **(e)**. **f** Proportional distribution of treatment groups within RL_pro- and NSC-like tumor cells. Two-sided t-tests were used to compare the indicated groups. *$P < 0.05$, **$P < 1.0e-10$, ***$P < 1.0e-20$.

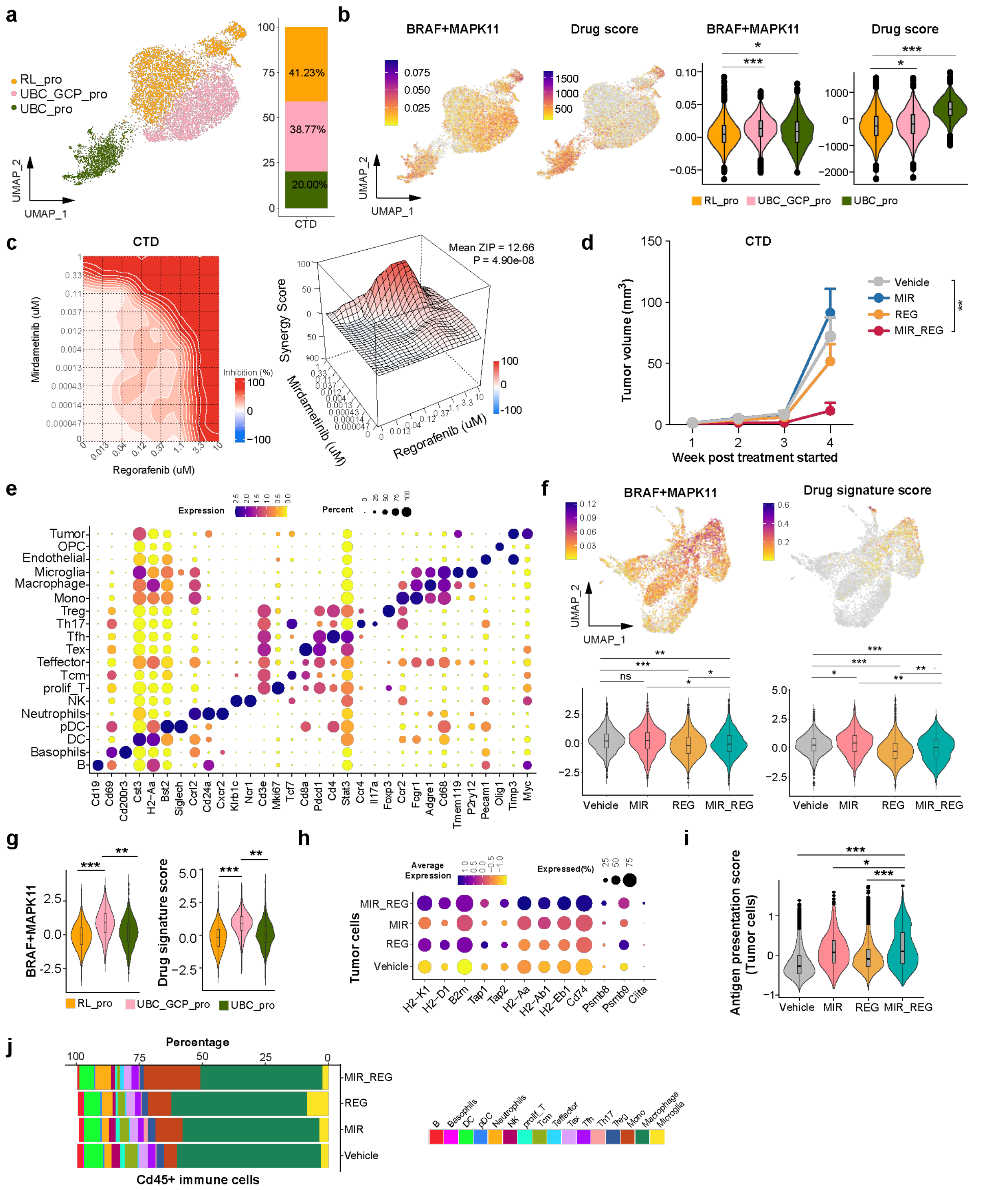


**Figure S10. Effects of combination therapy on CTD tumor cells and tumor microenvironment. a** Tumor cell states in CTD mapped using normal reference marker genes **Fig.5a**, with bar plot showing proportions of CTD cell states. **b** Feature and violin plots of driver combination activities and predicted drug sensitivity scores across tumor cell states in the CTD model. Two-sided t tests were used to compare RL_pro with UBC_GCP_pro and UBC_pro. *P < 0.05, **P < 1.0e-10, ***P < 1.0e-20. **c** Cell growth inhibition contour plot and synergy landscapes of regorafenib and mirdametinib combination treatment in CTD cells. **d** Changes in MRI-measured tumor volume in CTD-bearing mice after treatment initiation across different treatment arms. **e** Dot plot showing marker gene expression patterns in immune cell types and tumor cells. **f** Feature and violin plots of driver combination activities and predicted drug sensitivity scores across treatment arms from CTD preclinical study. **g** Violin plots of driver combination activities and predicted drug sensitivity scores across tumor cell states. **h** Dot plot showing antigen presentation gene expression in tumor cells across the four treatment groups. **i** Antigen presentation module scores in tumor cells across treatment groups; two-sided t tests were used to compare combination treatment with other groups. *P < 0.05, **P < 1.0e-10, ***P < 1.0e-20. **j** Proportional distribution of Cd45+ immune cells across treatment groups.

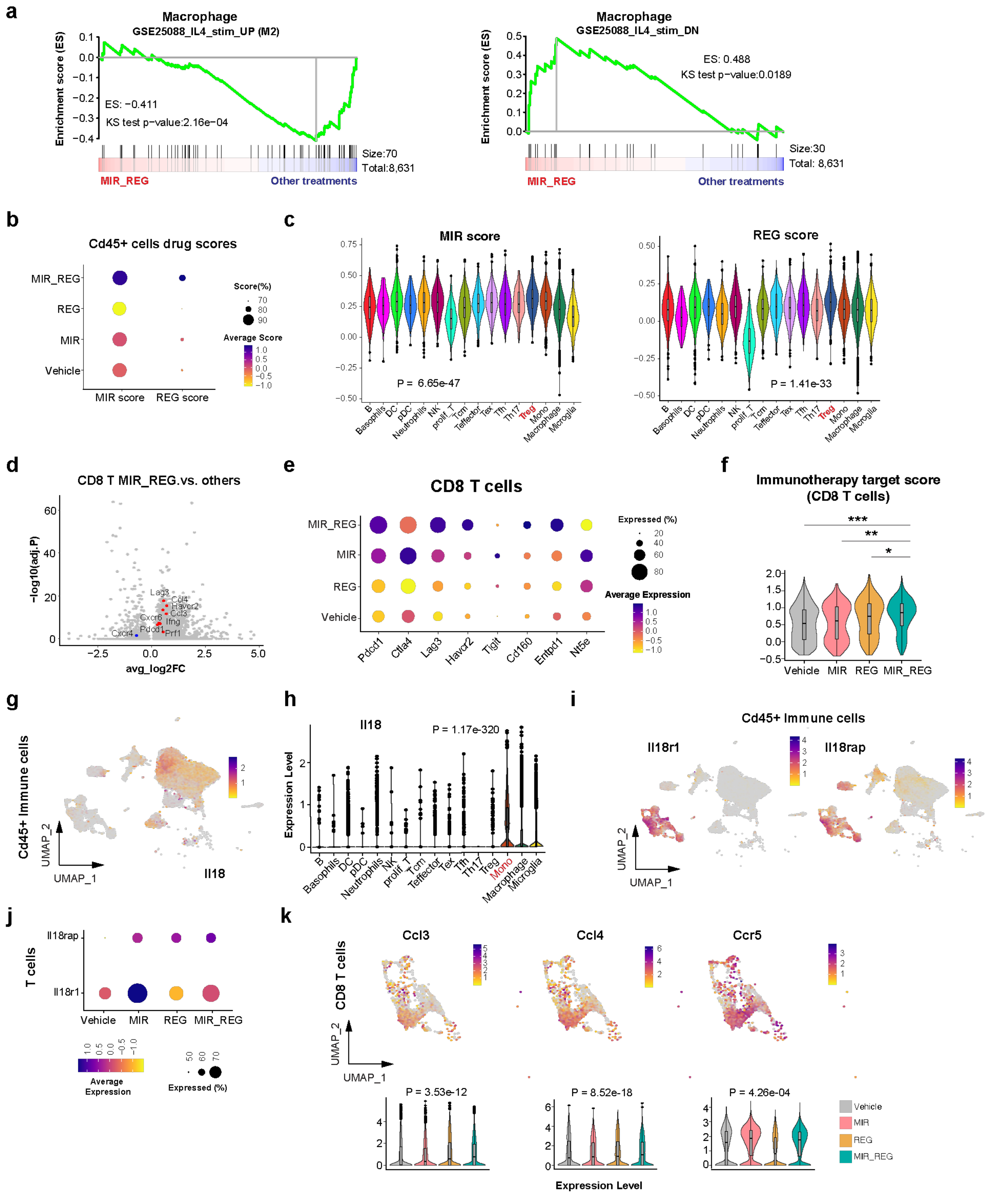


**Figure S11. The combination of MEK inhibition and regorafenib reconfigures the tumor microenvironment. a** Gene set enrichment analysis of M2-macrophage related genes in macrophages, comparing combination treatment versus other treatment groups. **b** Dot plot of mirdametinib and regorafenib drug signature scores in immune cells across four treatment groups. **c** Violin plot of mirdametinib and regorafenib drug signature scores across immune cell types; two-sided t test comparing Treg cells with other immune cells. **d** Volcano plot of differentially expressed genes in CD8+ T cells from the MIR_REG combination group versus other treatments. **e** Dot plot showing immune therapy–related target gene expression in CD8+ T cells across treatment groups. **f** Module scores for immune therapy target genes; two-sided t tests comparing combination treatment with other groups. *P < 0.05, **P < 1.0e-10, ***P < 1.0e-20. **g** Feature plot of Il18 expression in immune cells from the CTD model. **h** Violin plot of Il18 expression across immune cell types; two-sided t test comparing monocytes with other immune cells. **i** Feature plots of Il18r1 and Il18rap expression in immune cells from the CTD model. **j** Dot plot of Il18r1 and Il18rap expression in T cells across four treatment groups. **k** Feature and violin plots of Ccl3, Ccl4, and Ccr5 expression in CD8+ T cells across treatment groups; two-sided t test comparing combination treatment with other groups.

### Supplementary Tables

**Table S1. Unique genes, drugs, interactions, and drug phases in the six built-in drug-gene interaction databases, related to Figure 1.**

**Table S2. Bulk RNA-seq profiles from medulloblastoma patient cohorts, related to Figure 1, Supplementary Figure 1 and 8.**

**Table S3. The correlation between seed-driver activity or expression with MB patient survival, related to Supplementary Figure 1.**

**Table S4. BPdb included 658 BBB+ small molecular drugs, related to Figure 1.**

**Table S5. Seed drug and partner drug libraries for HTS, related to Figure 1 and Supplementary Figure 2.**

**Table S6. Two-dose combination screening, related to Figure 1 and Supplementary Figure 2.**

**Table S7. 10-dose combination screening, related to Figure 1.**

**Table S8. Normal glutamatergic lineage neuron developmental reference 10x scRNA-seq data QC, related to Figure 5, Supplementary Figure 7.**

**Table S9. MB patients sample information from the public single-cell RNA-seq datasets, related to Figure 5, Supplementary Figure 7, and 8.**

**Table S10. GSE155446 MB patient 10x scRNA-seq data QC, related to Figure 5, Supplementary Figure 7 and 8.**

**Table S11. Hovestadt et al. MB patient SMART-seq2 data QC, related to Supplementary Figure 7.**

**Table S12. Luo et al. MB patient 10x scRNA-seq data QC, related to Supplementary Figure 7.**

**Table S13. D341 and SJMB016880 *in vivo* 10x scRNA-seq data QC table, related to Figure 5, Supplementary Figure 9.**

**Table S14. CTD *in vivo* 10x scRNA-seq data QC table, related to Figure 7, Supplementary Figure 10 and 11.**

**Table S15. M1/G2 microphage signature gene sets, related to Supplementary Figure 11.**

**Table S16. Trametinib, regorafenib and mirdametinib drug signature genes sets, related to Figure 2, 4, 5 and Supplementary Figure 4, 8, 10 and 11.**
